# Non-oncology drugs are a source of previously unappreciated anti-cancer activity

**DOI:** 10.1101/730119

**Authors:** Steven M. Corsello, Rohith T. Nagari, Ryan D. Spangler, Jordan Rossen, Mustafa Kocak, Jordan G. Bryan, Ranad Humeidi, David Peck, Xiaoyun Wu, Andrew A. Tang, Vickie M. Wang, Samantha A. Bender, Evan Lemire, Rajiv Narayan, Philip Montgomery, Uri Ben-David, Yejia Chen, Matthew G. Rees, Nicholas J. Lyons, James M. McFarland, Bang T. Wong, Li Wang, Nancy Dumont, Patrick J. O’Hearn, Eric Stefan, John G. Doench, Heidi Greulich, Matthew Meyerson, Francisca Vazquez, Aravind Subramanian, Jennifer A. Roth, Joshua A. Bittker, Jesse S. Boehm, Christopher C. Mader, Aviad Tsherniak, Todd R. Golub

## Abstract

Anti-cancer uses of non-oncology drugs have been found on occasion, but such discoveries have been serendipitous and rare. We sought to create a public resource containing the growth inhibitory activity of 4,518 drugs tested across 578 human cancer cell lines. To accomplish this, we used PRISM, which involves drug treatment of molecularly barcoded cell lines in pools. Relative barcode abundance following treatment thus reflects cell line viability. We found that an unexpectedly large number of non-oncology drugs selectively inhibited subsets of cancer cell lines. Moreover, the killing activity of the majority of these drugs was predictable based on the molecular features of the cell lines. Follow-up of several of these compounds revealed novel mechanisms. For example, compounds that kill by inducing PDE3A-SLFN12 complex formation; vanadium-containing compounds whose killing is dependent on the sulfate transporter SLC26A2; the alcohol dependence drug disulfiram, which kills cells with low expression of metallothioneins; and the anti-inflammatory drug tepoxalin, whose killing is dependent on high expression of the multi-drug resistance gene ABCB1. These results illustrate the potential of the PRISM drug repurposing resource as a starting point for new oncology therapeutic development. The resource is available at https://depmap.org.

---

The prospect of repurposing existing drugs for new clinical indications is alluring: rapid clinical translation can, in principle, occur for drugs already proven safe in humans. To date, most oncology repurposing discoveries have been serendipitous; systematic, at-scale screening of the entire pharmacopeia has not been feasible.

Recent efforts, however, have demonstrated the power of large-scale cancer cell line screening – testing either a large number of compounds across a limited number of cell lines (e.g. the NCI-60 panel^1^), or a modest number of oncology compounds across a large number of cell lines (e.g. the GDSC project at the Sanger Institute^2^ and the CTD2 project at the Broad Institute^3^) (Fig. 1a). The ideal study would involve screening a large number of drugs (the majority of which are non-oncology drugs) across a large panel of genomically characterized cell lines, so as to capture the molecular diversity of human cancer. The extent to which non-oncology drugs have potential as future cancer therapeutics is unknown.

**Fig. 1.**
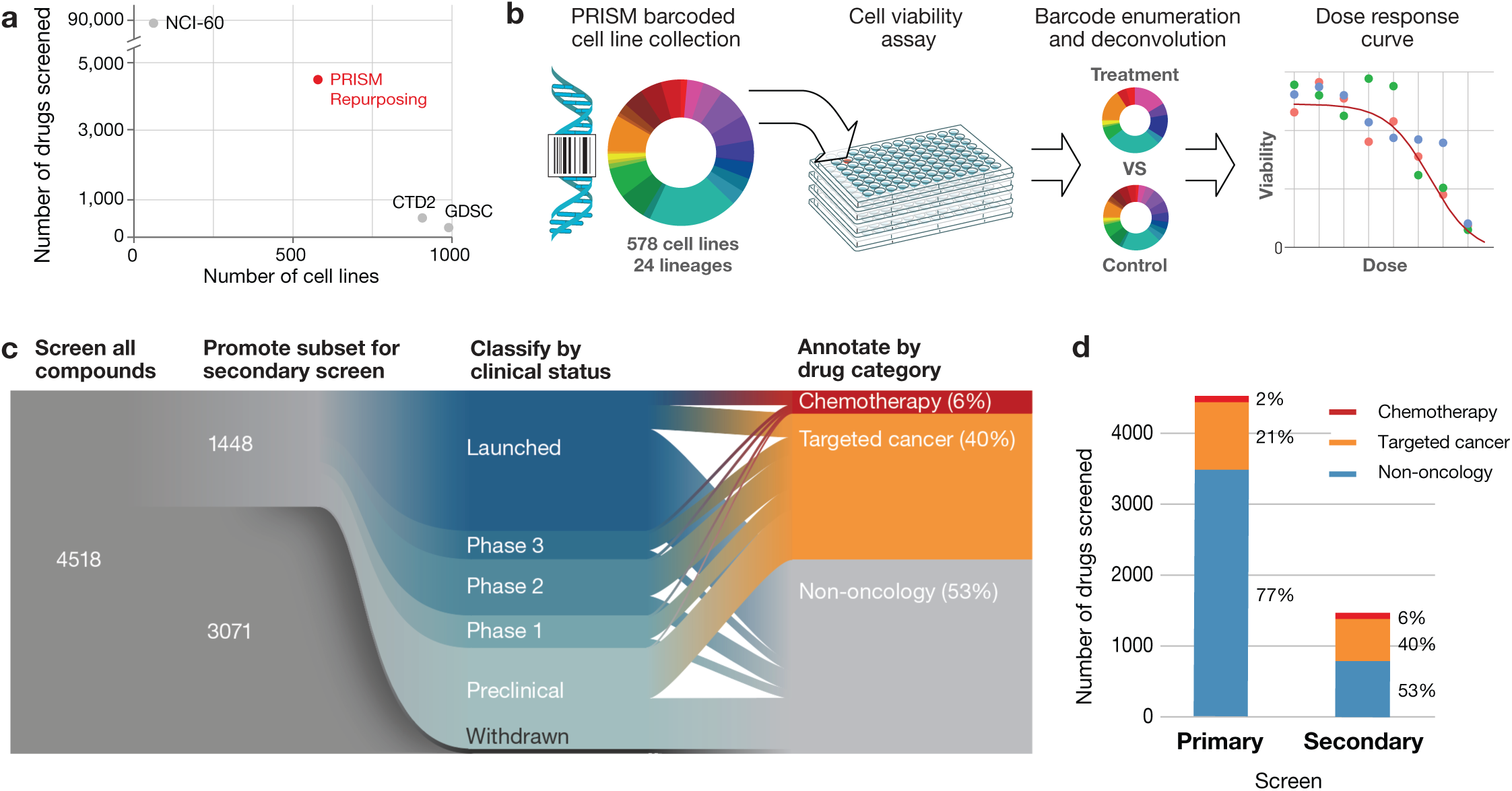
Generation of the PRISM Repurposing dataset. **a**, Dimensionality of publicly-available pharmacogenomic drug screening experiments. The PRISM Repurposing dataset contains approximately 10-fold more compounds than CTD2 and approximately 10-fold more cell lines than NCI-60. **b**, PRISM method overview. In the PRISM platform, barcoded cell lines are pooled in groups of 25 cell lines and treated with chemical perturbagens. Cell line pools are lysed and collapsed five days after perturbation and the abundance of mRNA barcodes present is measured using Luminex MagPlex Microspheres. Fluorescence values are deconvoluted to measure cell viability. **c**, Repurposing screen workflow. Primary screen of 4,518 drugs was performed at 2.5 μM. 1,448 active drugs were re-tested at 8 doses. Compounds were annotated as chemotherapy drugs, targeted cancer drugs or non-oncology drugs based on approved indications and prior clinical trial disease areas. **d**, Drug category representation in the primary and secondary screens. The secondary screen was enriched for chemotherapy and targeted cancer therapies.

We report here the feasibility of using the PRISM molecular barcoding and multiplexed screening method to test 4,518 existing drugs against 578 cancer cell lines. We find that non-oncology drugs have an unexpectedly high rate of anti-cancer activity, and we find that the sensitivity of cancer cell lines to many of these compounds can be predicted from the genomic features of the cell lines, thereby suggesting potentially relevant patient populations.

## RESULTS

### Drug selection and PRISM profiling

To facilitate the screening of thousands of compounds across hundreds of cell lines, we utilized the PRISM method, which involves the molecular barcoding of cancer cell lines with unique DNA sequences, thereby allowing the barcoded cell lines to be pooled with relative barcode abundance serving as a surrogate for cellular viability^4^ (Fig. 1b). Using PRISM, we screened 578 adherent cell lines spanning 24 tumor types (Extended Data Fig. 1a, Supplementary Table 1).

We chose 4,518 drugs from the Drug Repurposing Hub^5^ (https://www.broadinstitute.org/repurposing), and confirmed the identity and purity of all compounds by liquid chromatography-mass spectrometry (LC-MS) to be greater than 75% pure (Supplementary Table 2). 3,350 of the compounds (74%) are either approved for clinical use in the United States or Europe, or are in clinical development. The remaining 1,168 (26%) are tool compounds with known activities. The majority of the compounds, 3,466 (77%), were non-oncology related, with the remaining compounds being either chemotherapeutics (2%) or targeted oncology agents (21%).

### Screening results

We employed a 2-stage screening strategy whereby drugs were first screened in triplicate at a single dose (2.5 µM). 1,448 screening positives were then re-screened in triplicate in 8-point dose response ranging from 10 µM to 600 pM (Fig. 1c, Supplementary Table 2). Interestingly, the majority of active compounds (774/1,448, 53%) were originally developed for non-oncology clinical indications (Fig. 1d). The screening datasets are available on the Cancer Dependency Map portal at https://depmap.org and Figshare (doi:10.6084/m9.figshare.9393293) (Extended Data Fig. 1-7). We compared the PRISM results to two gold standard datasets: GDSC^2^ and CTD2^3^. The three datasets shared 84 compounds tested on a median of 236 common cell lines, yielding 16,650 shared data points. The PRISM dataset had a similar degree of concordance to GDSC and CTD2 (Pearson correlations over all shared data points 0.62 and 0.63, respectively), as the GDSC and CTD2 datasets had to each other (Pearson correlation 0.60) (Extended Data Fig. 8a). The three datasets were similarly concordant when the analysis was restricted to compounds showing evidence of anti-cancer activity (Extended Data Fig. 8b). We conclude that despite differences in assay format, sources of compounds^5^, and sources of cell lines^6^, the PRISM Repurposing dataset is similarly robust compared to other pharmacogenomic datasets.

We do note, however, that at the level of individual compound dose responses, the PRISM Repurposing screen tends to be somewhat noisier with a higher standard error in vehicle control measurement (Extended Data Fig. 8c, Extended Data Fig. 9). This variation may be explained by a combination of a longer assay duration, a smaller number of cells assayed, and/or variation attributable to growing cells in pools. Such noise, however, was not substantial enough to preclude the discovery of anti-cancer activities or their associated predictive biomarkers (see below).

### Landscape of non-oncology drug effects on cancer viability

We performed unsupervised clustering of compound viability profiles independent of their functional annotations using the Uniform Manifold Approximation and Projection (UMAP) method^7^ (Fig. 2a). Compounds with similar mechanisms of action (MOA) tended to cluster together, indicating that expected activities were recovered by the PRISM assay. Interestingly, while we expected to recover known MOAs for cancer drugs, we also found clusters of functionally-related *non-cancer* drugs, such as vitamin D receptor agonists and HMG-CoA reductase inhibitors. Of note, some functionally-related classes (e.g. glucocorticoid receptor agonists) showed two or more distinct clusters, suggesting that biologically-relevant substructure may exist within the dataset.

**Fig. 2.**
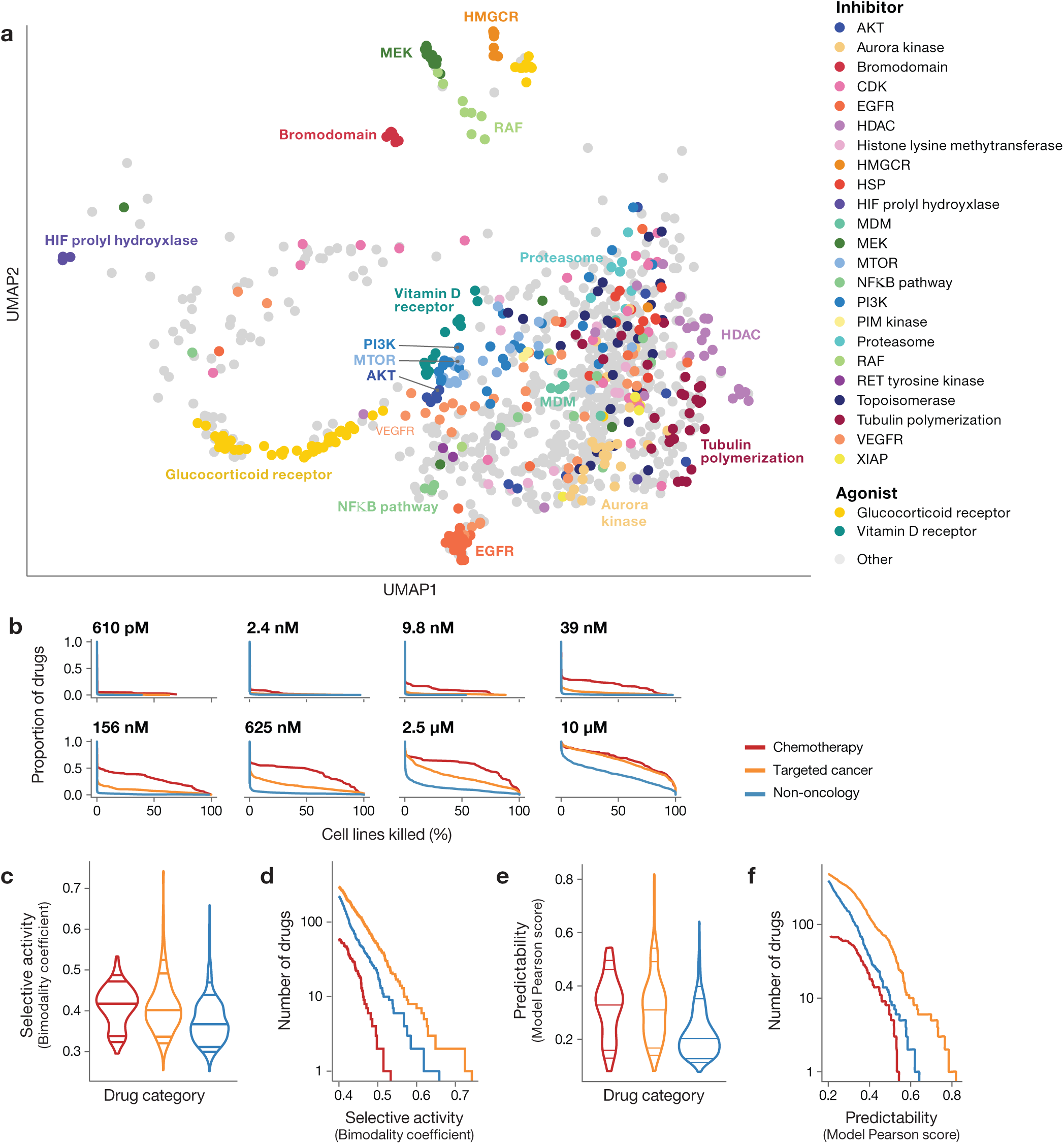
Drug response landscape of human cancer cell lines. **a**, Two-dimensional UMAP projection of drug killing profiles by cosine similarity. Clusters of compounds with shared existing mechanism of action are labeled by color. Compounds with average replicate Pearson correlation below 0.25 are not shown. **b**, Secondary screen drug activity by dose level. Dose-responsive killing is observed across all drug categories (chemotherapy: n = 90, targeted cancer: n = 584, non-oncology: n = 584) as indicated by color. Complementary cumulative distribution functions for the percent of cell lines killed in each drug category at each dose of the secondary screen is shown. **c**, Selective compound activity by drug category. Global distribution of secondary screen bimodality coefficients is shown. Dosewise bimodality coefficients are calculated from log fold-change level data. The maximum of all dosewise bimodality coefficients is shown for each compound. **d**, Most selective sensitivity profiles by drug category. The number of drugs (y-axis) from each drug category with a bimodality coefficient at any dose greater than a given threshold (x-axis). For visualization purposes, only drugs with bimodality coefficient greater than 0.4 are included. **e**, Predictability of compound activity by drug category. Global distribution of secondary screen Pearson scores is shown. ATLANTIS random forest models are trained to predict PRISM log fold-change scores using cell line baseline omics and genomic perturbation profiles. Pearson scores are defined as the correlation between actual and predicted PRISM profiles. The maximum Pearson scores across all models is shown for each compound. **f**, Most predictable sensitivity profiles by drug category. The number of drugs (y-axis) from each drug category with a predictive model with a Pearson score greater than a given threshold (x-axis). For visualization purposes, only drugs with a maximum Pearson score greater than 0.2 are included.

In general, chemotherapeutics killed the most cell lines, and non-oncology drugs the fewest, with targeted oncology drugs being intermediate (Fig. 2b). However, this pattern was highly dose-dependent: at high doses, targeted agents lost their selectivity. Perhaps most interestingly, a subset of non-oncology drugs showed particularly potent activity: 91 drugs killed at least 1% of cell lines at a concentration of 625 nM or lower.

To further investigate the therapeutic potential of non-oncology drugs, we computed the bimodality coefficient^8^ of each compound’s dose-wise viability profile (Extended Data Fig. 10a, Supplementary Table 3) and then calculated the maximum for each compound. While non-oncology drugs showed less bimodality than cancer drugs on average (Fig. 2c), the most selective compounds in the dataset include non-oncology drugs (Fig. 2d). This provides further evidence that large-scale testing of non-cancer drugs can reveal selective anti-cancer activity.

### Predictive models of killing activity

We next addressed the extent to which cell-killing activity was predictable based on the genomic features of the cell lines. For each drug, we used the random forest-based ATLANTIS algorithm^9^ using the baseline molecular features (cell lineage; gene copy number; function-damaging, hotspot, or missense mutations; DNA methylation levels; mRNA, miRNA, protein or metabolite abundance) and genetic dependencies (genome-scale CRISPR/Cas9 knockout or RNA interference) as inputs to the model^10–14^. Interestingly, mRNA expression was by far the most predictive feature type compared to other types of genomic information (Extended Data Fig. 10b).

The majority of highly predictable killing profiles came from targeted oncology drugs, compared to chemotherapeutics or non-oncology drugs (Fig. 2e, Supplementary Table 4-6). This is not surprising, as targeted oncology drugs have been optimized for their selective killing of subtypes of cancer. More striking was the observation that a substantial number of non-oncology drugs had highly predictable patterns of killing (Fig. 2f). In the primary screen, 195 non-oncology drugs had predictable killing with Pearson score > 0.2, and 23 were predictable with a Pearson score > 0.4. This same phenomenon of highly predictable non-oncology drug killing was observed when predictability was plotted against the bimodality score (Fig. 3). Importantly, only rarely (in 1% of cases) did the pattern of killing by non-oncology drugs correlate with knockout or knockdown of the drug’s intended target (Extended Data Fig. 10c, Supplementary Table 7). This suggests that the unexpected anti-cancer activity of non-oncology drugs is most likely to be explained by a previously unrecognized mechanism of action.

**Fig. 3.**
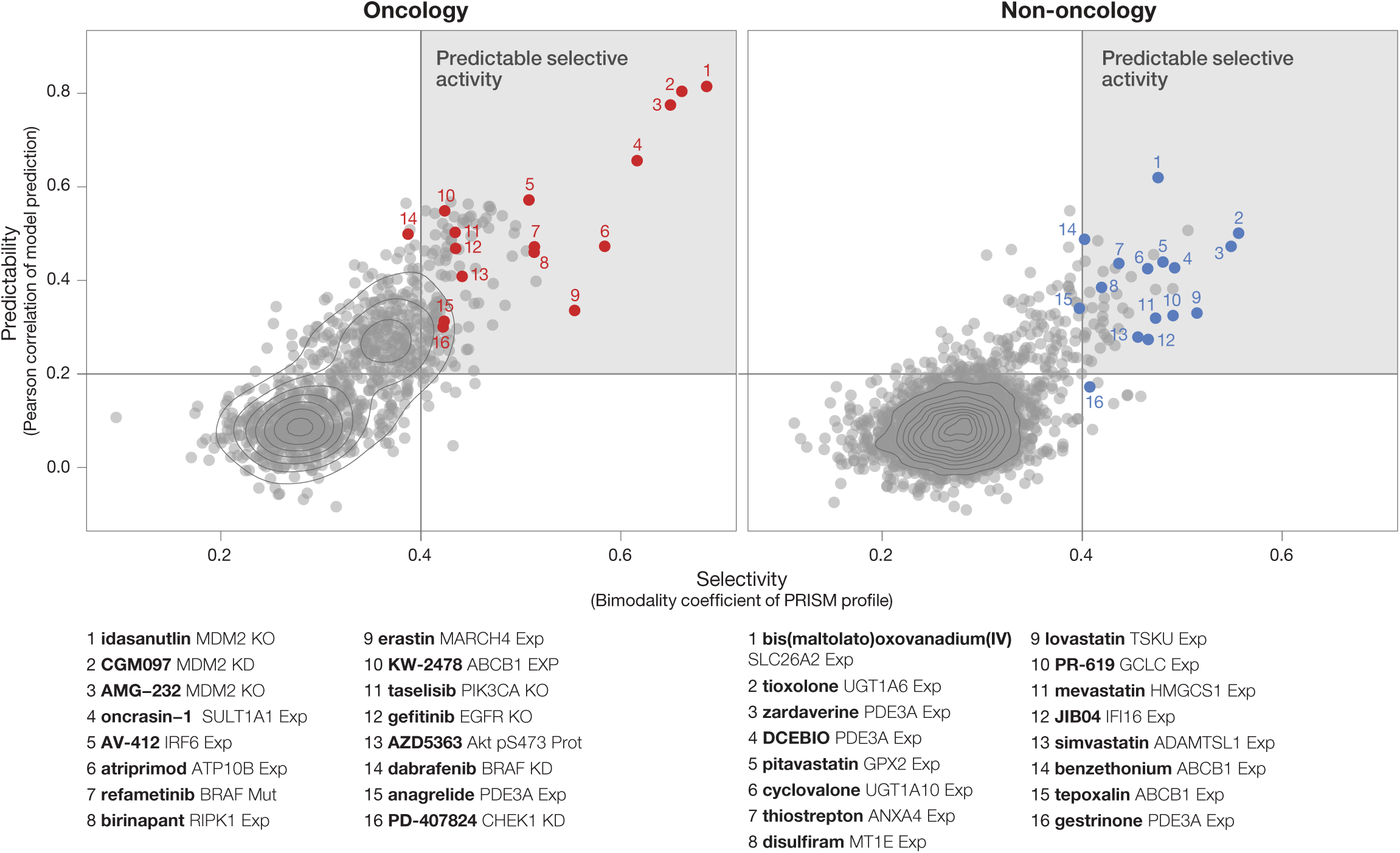
Predictive biomarker discovery using the PRISM Repurposing dataset. Prioritization of compounds based on strength of predictive models (Pearson score) and cell line selectivity (bimodality coefficient). Both known oncology and non-oncology drug sets yield hits with high selectivity and strong model performance. Compounds with highly predictable selective activity are shown in the upper right quadrant.

### Inducers of PDE3A-SLFN12 protein-protein interaction

Among the genomic features most highly correlated with non-oncology drug activity was expression of the phosphodiesterase gene PDE3A, whose expression correlated with killing by 11 structurally diverse compounds. These included the known PDE3A inhibitors anagrelide and zardaverine, progesterone receptor agonists (including the nonsteroidal drug tanaproget), the kinase inhibitor AG-1296 and the potassium channel activator DCEBIO (Fig. 4a). This pattern of killing was of particular interest because of the recent report of cancer cytotoxicity occurring as a result of protein-protein interaction between PDE3A and the largely uncharacterized protein SLFN12^15^. We found that the structurally diverse compounds identified in the PRISM screen bound PDE3A in a thermal shift assay (Fig. 4b, Supplementary Table 8), inhibited PDE3A enzymatic activity (Supplementary Table 9), and that their cytotoxicity was completely rescuable by either PDE3A knockout or by competition with trequinsin, a potent PDE3A small-molecule inhibitor that does not induce PDE3A-SLFN12 interaction^15^ (Fig. 4c). Importantly, PDE3A pull-down resulted in co-immunoprecipitation of V5-tagged SLFN12 following compound treatment, indicating that these novel compounds indeed induce PDE3A-SLFN12 protein-protein interaction (Fig. 4d). Taken together, the PRISM results show that an unexpectedly large number of structurally diverse non-oncology drugs kill PDE3A-expressing cancer cells by stabilizing the PDE3A-SLFN12 interaction. While these new compounds have limited potency, they may prove useful for starting points for further medicinal chemistry optimization of this novel anti-cancer mechanism.

**Fig. 4.**
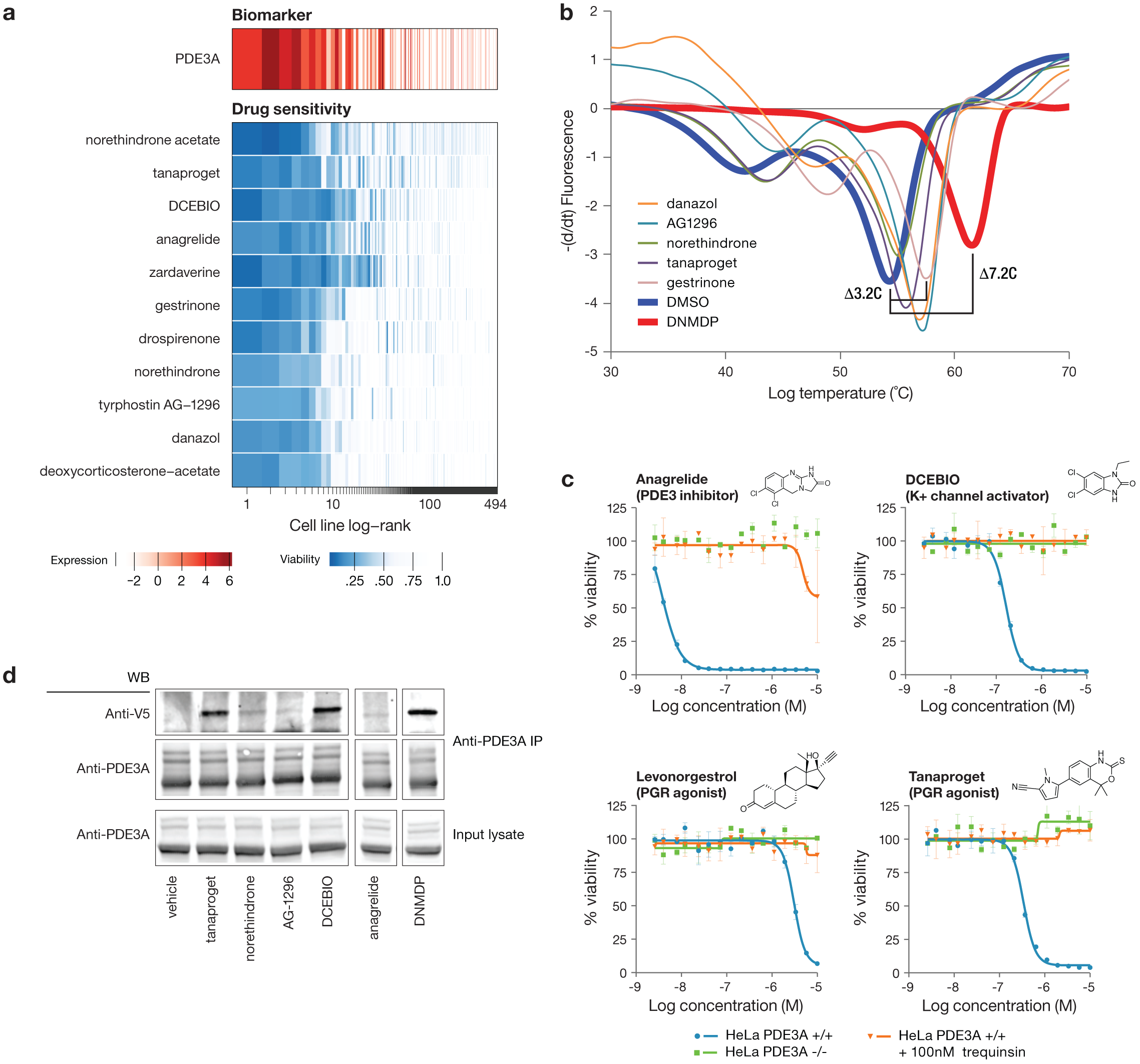
Multiple existing drugs selectively kill cancer cell lines by stimulating PDE3A-SLFN12 interaction. **a**, Drug sensitivity profiles associated with high PDE3A RNA expression. Compounds were selected from the PRISM 2.5 μM primary screen where PDE3A expression was the top predictive biomarker by ATLANTIS. Cell line viability is depicted versus PDE3A gene expression from CCLE RNAseq log2 RPKM data. Cell lines are ranked by mean viability of indicated compounds with a ceiling at 100%. **b**, Thermal shift assay performed in vitro with recombinant PDE3A. Results demonstrate stabilization of PDE3A protein for a subset of hit compounds. DNMDP is included as established positive control for strong PDE3A binding. Full ΔTm results are shown in Table S9. **c**, HeLa cell line dose response curves with PDE3A genetic loss or pharmacologic inhibition. Knockout of PDE3A was performed by CRISPR/Cas9. Parental HeLa cell killing was rescued by co-treatment with 100 nM of the non-cytotoxic PDE3A inhibitor trequinsin. Standard deviation across two replicates is shown with error bars. **d**, Co-immunoprecipitation of V5-tagged SLFN12. Newly identified PDE3A modulating compounds induced complex formation between PDE3A and SLFN12. Anti-V5 western blot is shown using lysates from HeLa cells treated with the indicated compound for 8 hours. The anagrelide and DNMDP lanes were run on the same gel as all other samples and were cropped for figure purposes.

### Predictive biomarkers of disulfiram activity

An intriguing observation was the association of killing by disulfiram (Antabuse), an inhibitor of acetaldehyde dehydrogenase used to treat alcohol dependence, and chromosome 16q copy number. Examination of the relevant region of 16q^16^ revealed that both copy number loss and low expression of the metallothionein genes *MT1E* and *MT2A* were correlated (Pearson correlations of 0.33 and 0.23, respectively, across 560 lines) with disulfiram-induced cell killing (Fig. 5a). Disulfiram has been previously suggested as an anti-cancer agent^17^, and at least one clinical trial has shown hints of efficacy in lung cancer when used in combination with chemotherapy^18^. In the absence of a predictive biomarker, the magnitude of clinical benefit did not warrant further clinical investigation.

**Fig. 5.**
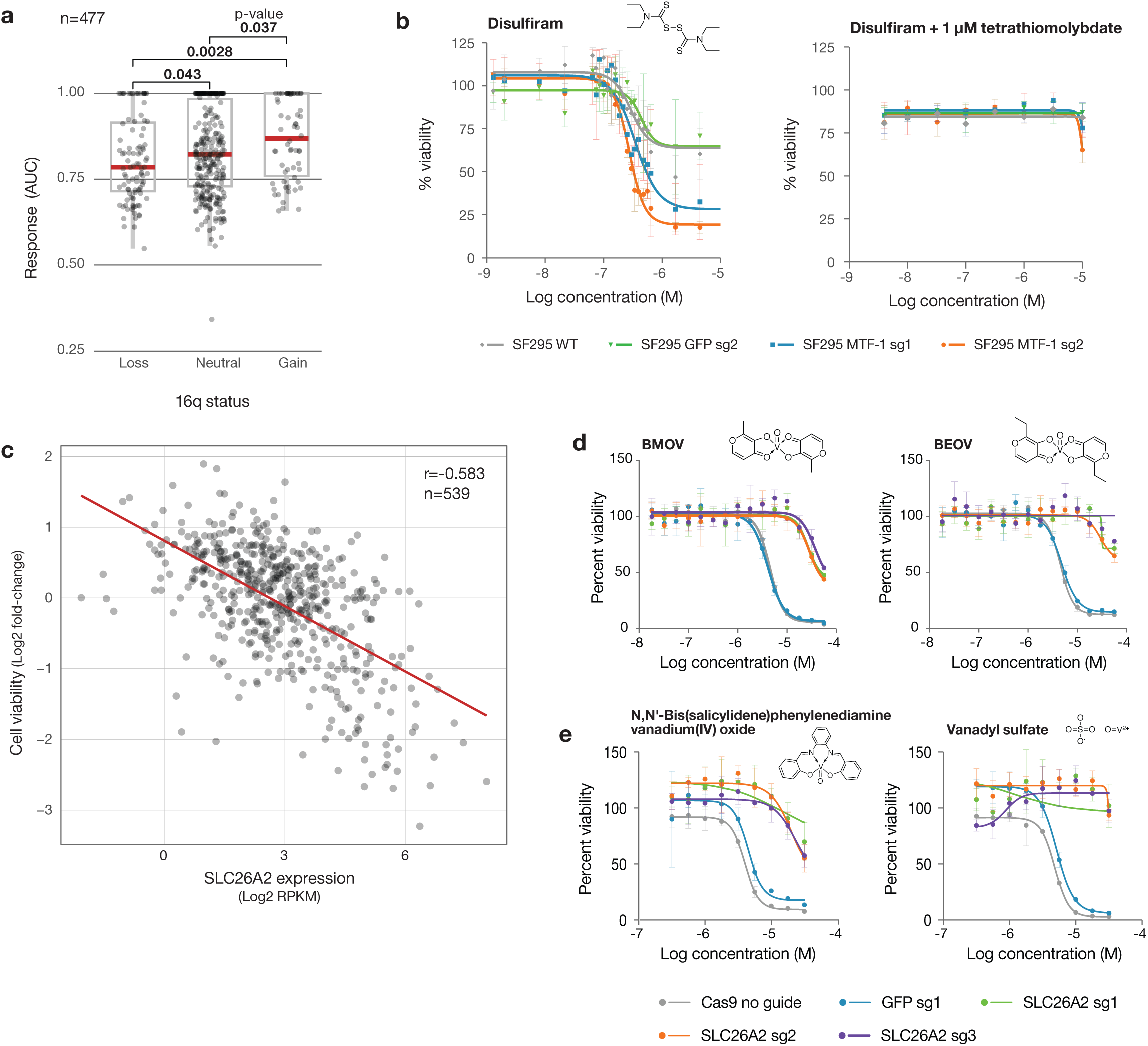
Anti-cancer activity of disulfiram and vanadium. **a**, Disulfiram sensitivity of PRISM cell lines grouped by 16q copy number status. PRISM secondary data shown as AUC. Arm-level copy number data was obtained using published TCGA methods and manually reviewed to ensure consistency with copy number at the 16q13 locus. Two-sided p-values were calculated using Wilcoxon signed-rank tests between each pair of groups. **b**, Drug sensitivity of SF295 cells with and without MTF-1 knockout. MTF-1 loss sensitizes to disulfiram. sgRNA against GFP was included as a non-targeting control. Co-treatment with 1 μM TTM rescues disulfiram killing in SF295 following MTF-1 knockout. Standard deviation across three replicates is shown with error bars. **c**, Scatter plot of BMOV sensitivity versus SLC26A2 gene expression. PRISM viability data at 2.5 μM is plotted against SLC26A2 gene expression from CCLE RNAseq (log2 RPKM). **d**, Dose-response curves for BMOV and BEOV in OVISE cells with and without CRISPR/Cas9-mediated knockout of SLC26A2. SLC26A2 knockout confers resistance. sgRNA against GFP was included as a non-targeting control. Standard deviation across 3 replicates is shown with error bars. **e**, Dose-response curves for related compounds, N,N’-Bis(salicylidene)-o-phenyldiamine vanadium(IV) oxide and vanadyl sulfate, in OVISE cells with and without SLC26A2 knockout. Standard deviation across 3 replicates is shown with error bars.

Our finding that the metallothionein genes *MT1E* and *MT2A* on 16q are predictive of disulfiram activity is mechanistically plausible, given that disulfiram’s activity is copper-dependent, and MT1E and MT2A are known metal-chelating proteins^19^. Consistent with this observation, *MT1E* and *MT2A* expression was also correlated with sensitivity to thiram and elesclomol, other copper-binding compounds^20,21^. In addition, disulfiram has been reported to induce metallothionein gene expression in prostate cancer cells^22^.

To test the hypothesis that metallothionein expression regulates disulfiram’s anti-cancer activity, we utilized the disulfiram-resistant glioma cell line SF295, which has an amplification of chromosome 16q and high metallothionein expression. In order to inhibit the expression of multiple metallothionein genes simultaneously, we knocked out the transcription factor MTF-1, which is a known upstream regulator of metallothionein gene expression (Extended Data Fig. 11a)^19^. Following MTF-1 knockout, metallothionein genes were among the most downregulated as assessed by global mRNAseq (Extended Data Fig. 11b). As predicted, MTF-1 knockout resulted in increased sensitivity of SF295 cells to disulfiram, and this increased sensitivity could be completely reversed by the copper chelator tetrathiomolybdate (TTM) (Fig. 5b). These results together suggest that 16q deletion is a potential predictive biomarker of disulfiram and other copper-dependent cytotoxic agents. This is particularly clinically relevant because arm-level 16q loss is seen in many tumor types, most notably breast and ovarian cancer, where its prevalence is estimated at 55-65% and 55-76%, respectively^23–25^.

### Vanadium-containing compounds

The PRISM screen revealed a strong correlation between killing by the vanadium-containing drug bis(maltolato)oxovanadium(IV) (BMOV) and expression of the sulfate transporter SLC26A2 (Pearson correlation −0.583, ATLANTIS Pearson score 0.620) (Fig. 5c). Bioavailable vanadium-containing compounds including BMOV and bis(ethylmaltolato)oxovanadium(IV) (BEOV) have been of interest for their ability to lower fasting blood glucose in animal models and patients with diabetes^26,27^. SLC26A2 function has not been extensively studied, but loss-of-function mutations have been associated with connective tissue disorders^28,29^. In cancer, it is broadly expressed at modest levels, with high expression in melanoma and uterine cancers^30^.

The mechanistic relationship between BMOV and SLC26A2 expression is not obvious. Arguing against BMOV being an inhibitor of SLC26A2 function, analysis of publicly available genome-wide CRISPR/Cas-9 loss-of-function screens indicates that SLC26A2 is not a cancer dependency (see https://depmap.org). Consistent with this, knockout of SLC26A2 in the BMOV-sensitive cell line OVISE resulted in no loss of viability (Extended Data Fig. 11). However, SLC26A2 knockout rendered OVISE cells resistant to both BMOV and BEOV, indicating that SLC26A2 is not simply a biomarker of killing, but is required for compound activity (Fig. 5d). The cytotoxicity of other, structurally-distinct vanadium-containing compounds was similarly rescued by SLC26A2 knockout, suggesting that the vanadium oxide ion is responsible for the SLC26A2-dependent cytotoxicity of BMOV (Fig. 5e). Whether these compounds are substrates for the SLC26A2 transporter, interfere with sulfate ion homeostasis, or confer some neo-function to SLC26A2 remains to be determined.

### Tepoxalin and multi-drug resistance

The expression of metabolic enzymes or drug efflux pumps were among the most common predictive biomarkers of drug response in the PRISM screen. As expected, high mRNA expression of the *ABCB1* transporter (encoding MDR1, also known as p-glycoprotein) was the top predictor of *resistance* to numerous oncology drugs including taxanes (docetaxel and paclitaxel), vinca alkaloids (vincristine and vinorelbine), and proteasome inhibitors (carlifizomib) (Fig. 6a).

**Fig. 6.**
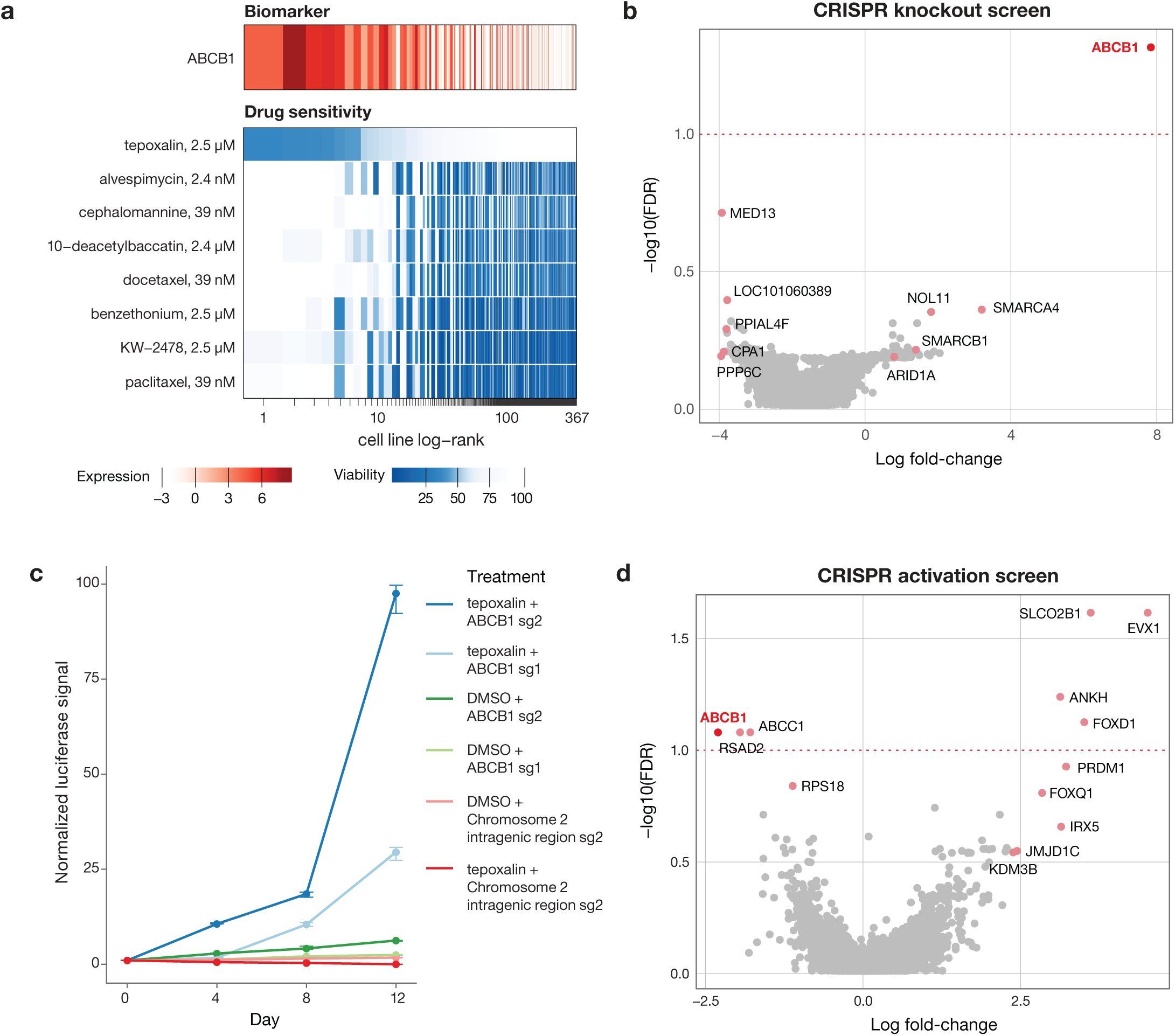
Tepoxalin is active against ABCB1-high cancer cell lines via an ABCB1-mediated mechanism. **a**, Drug sensitivity profiles of tepoxalin and seven additional compounds where ABCB1 RNA expression the predicted PRISM activity profile. Tepoxalin was the only compound tested where high ABCB1 expression was associated with sensitivity rather than resistance to tepoxalin. Cell line viability is depicted versus ABCB1 gene expression from CCLE RNAseq log2 RPKM data. Cell lines are ranked by mean viability with a ceiling at 100%. **b**, Analysis of tepoxalin genome-wide CRISPR/Cas9 gene knockout screen. ABCB1 knockout was top hit that rescued tepoxalin activity. LS1034-Cas9 cells (pXPR_311) were infected with the Brunello sgRNA library, selected with puromycin, and treated with 16 µM tepoxalin. Cells were passaged every 3-4 days over a 30-day period. Gene-level significance by MAGeCK-MLE versus log fold-change is shown. **c**, Tepoxalin cellular competition assay following ABCB1 knockout. LS1034-Cas9 cells (pXPR_311) were stably infected with firefly luciferase and parental LS1034 cells (without Cas9) were stably infected with renilla luciferase. Cells were mixed in 1:1 ratio an infected with the indicated sgRNA construct against ABCB1 or an intergenic region on chromosome 2 (negative control). Following puromycin selection, cell mixtures were treated with 16 µM tepoxalin. Firefly to renilla luminescence ratio is plotted as log fold-change over time. **d**, Analysis of tepoxalin genome-wide CRISPR/dCas9 gene activation screen. ABCB1 overexpression sensitizes to tepoxalin activity. LS1034-dCas9 (pXPR_109) were infected with the Calabrese sgRNA library, selected with puromycin, and treated with 16 µM tepoxalin. Cells were passaged every 3-4 days over a 14-day period. Gene-level significance by MAGeCK-MLE versus log fold-change is shown.

An unexpected finding, however, was that a single drug, tepoxalin, had the opposite relationship to ABCB1: high *ABCB1* expression predicted *sensitivity* to tepoxalin. Tepoxalin is a dual cyclooxygenase and 5-lipoxygenase inhibitor that is FDA-approved for the treatment of osteoarthritis in dogs^31,32^. Because more than 100 other cyclooxygenase and/or 5-lipoxygenase inhibitors were also tested in our PRISM screen, we asked whether they shared the ABCB1-associated killing effect. Strikingly, none of them did, suggesting that tepoxalin’s killing was most likely explained by an off-target mechanism. Consistent with this hypothesis, tepoxalin’s killing activity was not correlated with genetic knockout profiles of its known targets *PTGS1, PTGS2*, and *ALOX5*.

In order to gain insight into the mechanism by which tepoxalin selectively inhibits cancer cells, we performed a genome-wide CRISPR-Cas9 modifier screen to identify genes required for tepoxalin-mediated activity. LS1034 colorectal cancer cells have high levels of expression of *ABCB1* and are inhibited by tepoxalin with an IC50 of 3.8 µM. Cas9-expressing LS1034 cells were infected with a pooled library containing 76,441 sgRNAs targeting 19,114 genes and treated with 32 µM tepoxalin (or vehicle control) for 28 days. Remarkably, the gene knockout most enriched in tepoxalin-resistant cells was *ABCB1* itself (Fig. 6b). Other resistance hits included multiple components of the SWI/SNF complex (*SMARCA4, SMARCB1*, and *ARID1A*), which has been previously implicated in the regulation of *ABCB1* gene expression^33^. Consistent with these findings, cellular competition and dose response assays revealed strong selection for tepoxalin-treated ABCB1-null cells compared to wild-type cells (Fig. 6c, Extended Data Fig. 12a-c, Supplementary Table 10).

To complement the CRISPR-Cas9 loss-of-function screen, we performed a genome-wide CRISPR activation (CRISPRa) screen to identify genes whose overexpression was selected against in the setting of tepoxalin treatment. The gene showing the most negative selection was also *ABCB1*, indicating that its overexpression sensitizes cells to tepoxalin (Fig. 6d). In confirmatory studies, overexpression of ABCB1 in a low-expressing, tepoxalin-insensitive cell line resulted in increased drug sensitivity (Extended Data Fig. 12d-e). Furthermore, potent ABCB1 small molecule inhibitors did not phenocopy tepoxalin in the PRISM assay, and in fact antagonized tepoxalin-induced killing, indicating that ABCB1 inhibition alone does not explain tepoxalin-induced inhibition (Extended Data Fig. 12f-g).

Given that *ABCB1* encodes a drug transporter, we next asked whether ABCB1 expression affected intracellular concentrations of tepoxalin. Liquid chromatography-mass spectrometry (LC-MS) experiments indicated, however, that intracellular concentrations of tepoxalin were unaffected by levels of ABCB1 expression or by ABCB1 small-molecule inhibition (Extended Data Fig. 13a). Tepoxalin is also known to be metabolized to a compound known as RWJ20142 by conversion of its hydroxamic acid to a carboxylic acid (Extended Data Fig. 13b)^34^. Unlike tepoxalin, RWJ20142 was not cell permeable and showed no inhibition of ABCB1-expressing cancer cells. While tepoxalin did inhibit ABCB1 at high concentrations (Extended Data Fig. 13c-d), these results suggest that tepoxalin does not kill cancer cells by inhibiting ABCB1 activity, but rather that it requires ABCB1 for its cytotoxicity. Taken together, these results suggest that tepoxalin, but not its metabolite RWJ20142, inhibits *ABCB1* high-expressing cancer cells via a novel ABCB1-mediated mechanism.

### Data availability and analytical tools

The PRISM Repurposing dataset, including screening data and all metadata are available at the Cancer Dependency Map portal at https://depmap.org.

## DISCUSSION

The PRISM drug repurposing dataset described here represents to our knowledge the first-ever large-scale resource of anti-cancer activity of non-oncology drugs. The PRISM screen recovered 49 non-oncology compounds with selective and predictive biomarker-associated anti-cancer activity (Pearson score > 0.2, bimodality coefficient > 0.4) and 103 with a less stringent bimodality coefficient cutoff of > 0.35. Of note, 6 non-oncology compounds (Fig. 3) showed selective (bimodal) killing patterns, but their activity was found to be unpredictable based on the baseline genomic features of the cell lines. It is possible that an expansion of the number of cell lines in the PRISM panel will help to identify such biomarkers. Alternatively, killing might be explained by molecular features not yet measured in the cell lines.

It is conceivable that some non-oncology drugs could be brought directly to clinical trial for testing in cancer patients. However, before doing so, it will be important to establish that the killing activity of such drugs is observed at concentrations that are achievable and tolerable in humans. Similarly, it will be important to confirm that the predictive biomarkers identified in cell lines represent distinct populations of human tumors *in vivo*.

In contrast to repositioning of existing drugs for new indications, the PRISM results reported here also represent starting points for new drug development. In particular, when the anti-cancer activity of a drug occurs via an off-target mechanism, it is likely that further optimization for this new target will result in more potent and selective drug candidates. We note that the use of cell-based screens such as PRISM allows for the discovery of novel mechanisms of action that would be difficult to discover using conventional biochemical screening assays.

We initiated follow-up studies on 4 of the initial findings from the PRISM screen. In the case of disulfiram, we discovered a previously unrecognized biomarker (16q deletion) that predicts sensitivity. Future work will require extending such studies to the *in vivo* setting, and determining whether sufficiently high disulfiram concentrations can be achieved to obtain anti-cancer effects. The accessibility of copper in different organs and cell types will likely also modulate disulfiram’s anti-cancer activity. Our finding that compounds that kill PDE3A high-expressing cancer cell lines was striking because of the large degree of chemical diversity among these compounds. They all induced PDE3A-SLFN12 complex formation, as has been described for the compound DNMDP^15^. A structural understanding of the interaction of these diverse compounds with PDE3A is likely to inform future optimization of PDE3A-SLFN12-directed cancer therapeutics. Our finding that vanadium-containing compounds selectively kill cancer cells expressing high levels of the sulfate transporter SLC26A2 was also surprising, given that a mechanistic link between the two had not been previously suspected. A recent study showed that SLC26A2 expression is a mechanism of resistance to TRAIL-induced cell death^35^, but whether that mechanism is relevant to vanadium-induced killing, and whether these compounds dysregulate sulfate homeostasis remains to be determined. Perhaps most interesting was our observation that the drug tepoxalin has the unique ability to inhibit cells that express high levels of the multi-drug resistance gene *ABCB1*. While tepoxalin was originally developed as a cyclooxygenase/5-lipoxygenase inhibitor, our structure-activity relationship studies clearly showed that tepoxalin’s anti-cancer activity is likely cyclooxygenase and 5-lipoxygenase independent. While we showed that ABCB1 is both necessary and sufficient to confer tepoxalin cytotoxicity, the precise mechanism by which such cell death occurs remains to be established. Further optimization of tepoxalin against this new target, and engineering out its cyclooxygenase/5-lipoxygenase activity would likely result in improved tolerability as an anti-cancer agent.

We note that a challenge with cell-based phenotypic screens has historically been the difficulty in gaining molecular insight into hit compounds’ mechanism of action. Our results in the present study, however, demonstrate the power of genome-scale CRISPR/Cas-9 loss-of-function and gain-of-function screens to provide mechanistic clues to small-molecule action. For example, both CRISPR knockout and CRISPR activation screens pointed to ABCB1 as the most relevant target of tepoxalin. The availability of such functional genomic screening methods will likely reinvigorate cell-based screening generally. We also note that whereas small molecules are typically thought of as inhibitors of their protein targets, our PRISM results indicate that this is often not the case. For example, we discovered compounds that stabilize protein-protein interaction (e.g. PDE3A-SLFN12) and that likely engage ABCB1 but do not kill cells by ABCB1 inhibition. It is likely that a plethora of non-inhibitory small molecule activities remain to be discovered from this PRISM dataset.

The PRISM barcoding and pooling approach described here substantially increases screening efficiency, but it is conceivable that the pooling of cell lines results in paracrine-mediated mechanisms that modulate drug sensitivity. In practice, however, we have yet to observe such cell-cell interactions or any consistent discordance between PRISM and one-by-one viability profiling. Nevertheless, we and others have reported the existence of microenvironment-mediated drug resistance mechanisms^36^ and so the potential for such interactions should be considered when interpreting PRISM results.

The PRISM Repurposing dataset described here represents nearly half of all drugs ever tested in humans. Given the large number of unexpected findings that emerged from this initial screen, we believe that expansion of the PRISM resource in both the dimension of drugs and cancer models is warranted. Such data will provide an important pharmacological component of the Cancer Dependency Map (https://depmap.org), which in turn will form a pre-clinical foundation for cancer precision medicine.

## ACKNOWLEDGMENTS

We thank C. Yu, W. Hahn, B. Wolpin, A. Bass, N. Gray, E. Stover, T. Lewis, M. Mesleh, A. Burgin, S. Alper, G. Botta, M. Macaluso, P. Tsvetkov, X. Jin, K. Blakeslee, G. Ciolek, and E. Lander for helpful scientific discussions. M. Passino, C. Zhu, K. Gore, M. Laird, C. Trapechio, and E. Parikh generated barcoded PRISM cell lines and performed assays. Karolina Stumbraite assisted with STR fingerprinting. S. Johnson and J. Davis performed lysate processing and detection. S.E. Johnson and R. Singh provided analytical chemistry support. A. Vrcic, C. Sandland, and S. Figueroa-Lazu assisted with compound management. G. Kugener and A. Gonzalez provided technical assistance. Supported in part by the Carlos Slim Foundation (Slim Initiative in Genomic Medicine for the Americas (SIGMA)), NIH grants U01 HG008699 (T.R.G and A.S.), U54 HL127366 (T.R.G and A.S.), KL2 TR002542 (S.M.C.), and K08 CA230220 (S.M.C.), and the Conquer Cancer Foundation of ASCO Young Investigator Award (S.M.C.).

## AUTHOR CONTRIBUTIONS

Conceptualization, S.M.C., M.K., J.G.B., J.A.B., J.S.B., C.C.M., A.T., and T.R.G.; Methodology, S.M.C., R.T.N., R.D.S., M.K., J.G.B., D.P., E.L., R.N., J.A.B., C.C.M., A.T., and T.R.G.; Software, J.R., M.K., J.G.B., V.M.W., E.L., R.N., P.M., Y.C., M.G.R., and L.T.W.; Validation, S.M.C., R.T.N., R.D.S., J.R., M.K., and J.G.B.; Formal analysis, S.M.C., J.R., M.K., J.G.B., V.M.W., J.M.M., and L.T.W.; Investigation, S.M.C., R.T.N., R.D.S., J.R., M.K., J.G.B., R.H., D.P., X.W., S.A.B., N.J.L., U.B., N.D., P.J.O., A.S., and C.C.M.; Resources, E.S., J.G.D., H.G., M.M., F.V., A.S., J.A.R., J.A.B., A.T., and T.R.G.; Data Curation, S.M.C., R.D.S., and S.A.B.; Writing –Original draft, S.M.C., R.T.N., R.D.S., J.R., M.K., J.G.B., R.H., X.W., V.M.W., A.A.T., S.A.B., U.B., J.M.M., A.T., and T.R.G.; Writing – Review & Editing, M.G.R., and N.D.; Visualization, J.R., M.K., V.M.W., A.A.T., P.M., and B.T.W.; Supervision, M.M., A.T., and T.R.G.; Project Administration, S.A.B., A.S., J.A.R., and C.C.M.; Funding Acquisition, S.M.C., J.S.B., and T.R.G.

## METHODS

### PRISM screening

Parental cell lines were obtained from the Cancer Cell Line Encyclopedia (CCLE) project^10^ for PRISM cell line barcoding. This method was previously described^4^; however, we have made several improvements. The original paper described a 24-nucleotide barcode that was stably infected into cancer cell lines. The unique barcode identifier is located at the end of the blasticidin resistance gene to enable selection of infected cells. The barcode is also expressed as an mRNA under the highly active PGK promoter. This enabled us to detect the barcodes using a variant of the mRNA capture and Luminex detection method developed for the L1000 gene expression assay^37^, rather than by the previously described gDNA method. In addition to an mRNA versus gDNA readout, assay improvements include pooling cell lines according to doubling time similarity, increasing efficiency of screening by collapsing lysate plates by drug treatment, and the addition of a spike-in control for amplification and detection.

Barcoded cell lines were pooled (25 cell lines per pool) based on doubling time, then frozen into assay-ready vials. Vials were thawed and one pool was immediately plated per 384 well assay plate at 1,250 cells/well in triplicate. Cells were either treated the following morning with compounds by pin transfer (Repurposing primary and secondary HTS screens) or plated directly onto assay ready plates containing compounds (used for follow-up in the lower scale MTS004 and MTS006 screens). After a five-day incubation, cells were lysed. Lysate plates containing one pool of 25 cell lines each were then further pooled together to have one (in the secondary screen) or two (in the primary screen) final detection pools for amplification and barcode measurement. For the secondary screen, a set of ten unique barcodes were spiked-in to each well prior to PCR to control for variation in PCR amplification and Luminex detection following lysate pooling.

### Data processing

Luminex technology produces .lxb files containing fluorescence values for each Luminex bead observed during detection. Median Fluorescence Intensity (MFI) values were calculated as the median fluorescence values of all beads corresponding to a single PRISM barcode in a single technical replicate. MFI values are available in the “primary_MFI” and “secondary_MFI” files at https://depmap.org.

MFI values were log2-transformed (logMFI) and low-quality data was removed in a two-step process. First, an “outlier-pool filter” was applied. To detect probable screening artifacts, logMFI values were centered to the median logMFI for each cell line on each assay plate in order to put the measurements from each cell line on the same scale. For each well on each assay plate, the median of these centered values was computed. The assay-plate well medians were then standardized according to the median and median absolute deviation (MAD) across all assay-plate well medians from wells in the same position and assay plates associated with the same compound source plate. Data from wells with a standardized score of greater than 5 or less than −5 were excluded from all further processing steps.

Second, a “control separation filter” was applied. For each cell line on each plate, the distribution of logMFI values observed for the DMSO-treated negative controls was compared to that of the positive controls using a robust form of the Strictly Standardized Mean Difference (SSMD)^38^. Specifically, SSMD was calculated as:

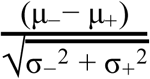

Where μ_-/+_ and σ_-/+_ stand for the medians and the MADs of logMFI values computed over the negative/positive control wells for each cell line on each plate. Data corresponding to SSMD values less than 2 were removed before calculating cell viability.

In the PRISM Repurposing primary screen, the outlier pool filter removed 2.4% of wells screened from further analysis (Extended Data Fig. 2). When applying the control separation filter, 488/578 cell lines (84%) had sufficient separation in more than 80% of assay plates (Extended Data Fig. 3). 497/578 cell lines (86%) had at least one passing replicate for every compound, and 377/578 cell lines (65%) had high-quality data for at least two replicates (Extended Data Fig. 4).

1,448 compounds were selected for secondary 8-point dose response testing based on reproducibility, predictability, and selectivity. In this particular PRISM screen, 15% of the assay plates had low quality, whereas the remaining 85% had data quality similar to the primary screen (92% of cell lines had at least one replicate that passed QC metrics, and 67% of cell lines had at least two passing replicates) (Extended Data Fig. 5-7). Additional improvements in assay automation have further improved data quality since this initial screen.

For the secondary screen only, ten unique barcodes were spiked into each well of each plate after cell lysis. Normalized MFI (nMFI) values were computed by taking the ratio of each logMFI value to the median logMFI of the inert barcodes in each well. For data produced before the spike-in protocol was introduced, nMFI values were set equal to MFI values.

Fold-change values were calculated as the ratio of nMFI to the median of the nMFI from the DMSO-treated negative controls for each cell line on each plate. Batch effects produced from variable detection and assay conditions were then removed using ComBat^39^. In particular, ComBat is run over each treatment condition separately by considering the log2-transformed fold-change values as probes and each pool-replicate combination as a batch (3 replicates x 20 pools = 60 batches). Corrected-log fold change values were then median collapsed for each cell line, screen, source plate, and well combination. We label cell lines as sensitive to a treatment if the median collapsed fold-change is less than 0.3.

The raw and processed data is available at https://depmap.org. Associated documentation and R scripts used for data processing will be available at https://github.com/broadinstitute/repurposing.

### Dose response

Measures of dose response were obtained by fitting 4-parameter logistic curves to viability values for each compound and cell line using the R package “drc”. Following the practice of Smirnov and Safikhani^40^, the upper asymptote of the logistic curves is fixed at 1 and the viability values are fit as a function of drug concentration according to:

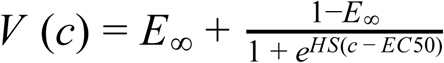

where all concentrations are in the natural logarithm scale. IC50 values were defined as the concentration c at which V(c) = 0.5. Additionally, the area under the dose response curve (AUC) was calculated using the normalized integral:

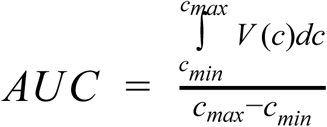

The formulation above puts AUC values on a scale between 0 and 1 for curves with lower asymptotes less than 1, where lower AUC values indicate increased sensitivity to the treatment.

### Biomarker discovery

To generate predictive biomarkers, we adopted ATLANTIS predictive models^9^ and trained multiple models for each PRISM profile. ATLANTIS is a tailored non-linear regression model for gene dependency prediction based on the baseline characteristics of cancer cell lines. More specifically, ATLANTIS is an efficient implementation of a conditional inference forest^41^ with additional weighting and iterative feature selection steps. Implementation details have been previously published and the code is available on a public repository (https://github.com/cancerdatasci/atlantis).

For each dose-wise log-fold change profile, 14 ATLANTIS models are trained (one model per feature set). Feature sets include baseline cell line omics, genetic dependencies and experimental confounders. All feature sets are listed in the Supplementary Table 11.

Next, the predictive performance of each model is assessed based on the Pearson correlation between out-of-bag model predictions and the response variable. Models with Pearson correlation larger than 0.2 are considered as strong models. Variable importance measures (“mean decrease in accuracy”) output by ATLANTIS are computed for each model. The most important feature of each strong predictive model is presented as a potential predictive biomarker or strongly associated phenotype. The comprehensive list of biomarkers is available at https://depmap.org.

### Compound Selectivity

To assess the selective killing activity of compounds, we computed the bimodality coefficient^8^ for each median collapsed log-fold change PRISM profile. The bimodality coefficient was defined as:

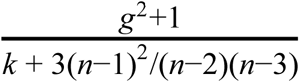

Where n is the number of samples (cell lines), g is the sample skewness and k is the sample excess kurtosis. Note that, larger bimodality coefficient implies a highly skewed (large magnitude of g) but light-tailed (small kurtosis) distribution.

### Nomination for Secondary Screen

Compounds from the primary screen were prioritized for secondary screening based on the existence of a strong biomarker, reproducibility of killing profile, selectivity of killing, and availability, and novelty of compound. 1,447 compounds were chosen among these prioritized compounds to be progressed to secondary profiling at 8-point dose.

### Computation of AUC values for Cross-Dataset Comparison

Secondary PRISM Repurposing data was compared to CTD2 (Version 2.0, December 15th 2015 version) and GDSC available through the PharmacoGX package^40,42^. For the PRISM Repurposing and GDSC datasets, dose response curves were fit as described above. CTD2 provides dose-response curve parameters and curves were not refit.

We limit the scope of the comparison to compounds screened in all three datasets. For each compound, the shared dose range across all three datasets was computed. A compound was included in the comparison if the maximum of the shared dose range was at least four times the minimum of the shared dose range (which guarantees at least one data point from all three datasets).

Dose response curves were computed for each compound-cell line-dataset combination using all available data, i.e. no dose range limitations. AUC values were then calculated over the shared dose range (curves were not refit). See the “Dose response” methods section for details.

The complete table of the published/recomputed dose response parameters and AUC values are given in Supplementary Table 12.

### Assessment of Noise in PRISM Repurposing Secondary Dataset

To quantify the amount of noise in the PRISM screen, we estimated the standard error of the inferred log fold-change viability values. In particular, we made the assumption that (spike-in) normalized logMFI values have independent additive zero mean errors with a constant variance σ^2^. A standard error propagation analysis (see section 9.3 of ^43^), gives

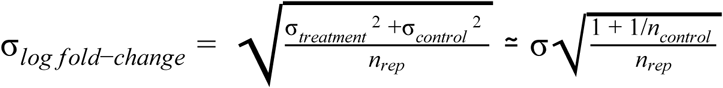

Where *n*_*rep*_ is the number of replicates (3 for PRISM Repurposing), *n*_*control*_ is the number of negative control wells in a given plate (32 in the standard PRISM assay format), and σ_*log fold*-*change*_ is the standard-error estimate.

We estimate the variance σ^2^ separately for each cell line, using three DMSO-only detection plates (compound plate PMTS009) in the MTS006 screen. In particular, we estimated the variance of the normalized log2 MFI values for each cell line under DMSO treatment on an assay plate basis. Next, we plug the median of assay plate variances into the formula above to obtain an estimate of σ _*log fold*-*change*_.

The same parameters are estimated both for the rest of the Repurposing library and also for the GDSC and CTD2 datasets (σ^2^ is estimated as the variance of the log2 intensity values in the DMSO after correcting for the background noise). The comparison of the noise levels through PRISM screens and other assays are presented in the Extended Data Fig. 9S.

### Projection of Viability Profiles to Two Dimensions

We used Uniform Manifold Approximation and Projection (UMAP)^7^ on the cosine similarity between unannotated compound log-viability (base 2) profiles from the primary screen to generate the two-dimensional projection in Figure 2A. From the primary screen compound viability matrix, we excluded cell lines that were missing more than 10% of their viability values (failed in QC steps) following compound treatment. The remaining missing values were imputed using the FastImputation R package^44^. We perform UMAP on the resulting matrix, using the cosine distance metric and the configuration parameters: 7 nearest neighbors, 0.5 minimum distance, 2 components, and 200 training epochs. The remainder of the parameters are set to the defaults provided by the UMAP package.

For the visualization, we filter out compounds that have an average replicate correlation between three replicates below 0.25. In addition, some MOAs that have fewer than 20 compounds are shown in translucent grey.

### Comparison to genetic loss-of-function screens

Linear models were fit to test the association between primary collapsed log fold-change profiles of each drug and DepMap Avana CRISPR knockout gene effect scores using the lmFit function from the limma R package with default parameters. For primary data, a single profile was selected to test for each drug. The procedure was repeated for secondary collapsed log fold-change profiles. A single dose series was selected for each drug.

### Antibodies and compounds for confirmatory studies

Paclitaxel (S1150), Tyrophostin AG 1296 (S8024), Anagrelide (S3172), Levonorgestrel (S1727), Deoxycorticosterone acetate (S4243), Drospirenone (S1377), and Norethindrone (S4040) were purchased from Selleck Chemicals. Dofequidar (SML0938-5MG), N,N′-Bis(salicylidene)-o-phenylenediamine vanadium(IV) oxide complex (68541), and vanadium(IV) oxide sulfate hydrate (233706) were purchased from Sigma-Aldrich. Tepoxalin (T103205) was purchased from Toronto Research Chemicals and WuXi AppTec (custom synthesis). DCEBIO (1422) and Zardaverine (1046) were purchased from Tocris. Disulfiram (HY-B0240), Bortezomib (HY-10227), and Tanaproget (HY-15606) were purchased from MedChemExpress. BMOV (FB18735) was purchased from Carbosynth. Tetrathiomolybdate (AC389530010) was purchased from Thermo Fisher Scientific. Danazol (1500220) was purchased from Microsource. Gestrinone (Prestw-1267) was purchased from Prestwick. The following antibodies were used: polyclonal rabbit anti-PDE3A from Bethyl Laboratory (A302-740A), monoclonal mouse anti-V5 from Life Technologies (R960-25), monoclonal rabbit anti-ABCB1 (D3H1Q) from Cell Signaling Technology (12683), monoclonal mouse anti-β-Actin (8H10D10) from Cell Signaling Technology, and monoclonal rabbit anti-SLC26A2 Antibody (3F6) from Novus Biologicals (H00001836-M04).

### Chemical synthesis

The compounds described below can be prepared from readily available starting materials using the following general methods and procedures.

### Synthesis of RWJ20142

Reactions were monitored by thin layer chromatography (TLC) with 0.25 mm E. Merck pre-coated silica gel plates (60 F_254_) and Waters Alliance HT LCMS system (Waters 2998 UV/Visible Detector, Waters Acquity SQD Mass, Waters e2795 Sample Manager) using Waters Cortecs C18 column (3 x 30 mm, 2.7 μm particle size). Additional parameters: solvent gradient = 97% A at 0min, 5% A at 1.75min, 97% A at 2.28 min, total 2.60 min; solvent A = Water (MilliQ) + 0.01% formic acid (Sigma); solvent B = Acetonitrile (EMD) + 0.01% formic acid (Sigma); flow rate: 1.75 mL/min. Purification of reaction products was carried out by flash chromatography using CombiFlash^®^Rf with Teledyne Isco RediSep^®^Rf High Performance Gold or Silicycle SiliaSep™ High Performance columns (4 g, 12 g, 24 g, 40 g, 80 g or 120 g. 1H NMR and 13C NMR spectra were obtained using a Bruker 400 Ascend. Chemical shifts are reported relative to chloroform (δ = 7.24) for ^1^H NMR.

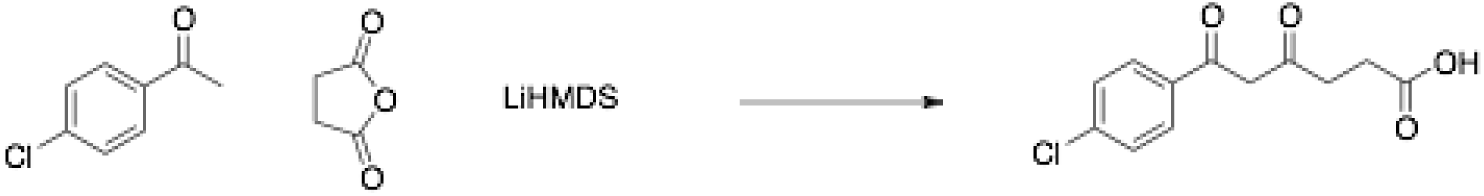

In a 100ml oven dried flask, a solution of LiHMDS (10 ml, 10 mmol, 1.0M in THF) was added dropwise to a solution of 1-(4-chlorophenyl)ethanone (1.10 g, 7.14 mmol) in dry THF (20 ml) at −78°C under argon atmosphere. After 1h, a solution of dihydrofuran-2,5-dione (0.86 g, 8.52 mmol) in dry THF (10 ml) was added at −78°C, stirred for 30 minutes then warmed up to room temperature. After 2h, the reacting mixture was quenched with water, acidified with 1N HCl (pH2), then extracted with DCM (3×10 ml). The combined organic layers were dried over Na_2_SO_4_, filtered and concentrated. The residue was purified by column chromatography on silica gel (10 to 50% AcOEt in hexanes) to afford 6-(4-chlorophenyl)-4,6-dioxohexanoic acid (550 mg, 31% yield). LCMS: MS(ESINeg) m/z = 253 [M-H]^-^.

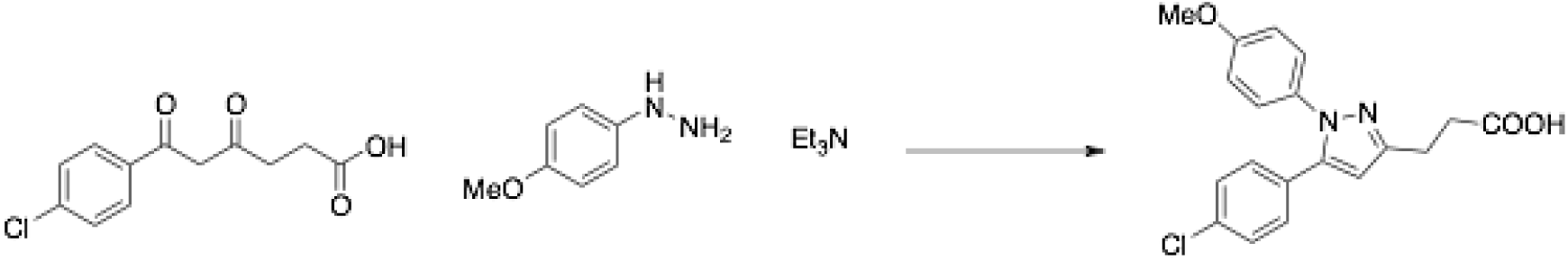

In a 50ml flask, a solution of (4-methoxyphenyl)hydrazine hydrochloride (0.37 g, 2.12 mmol), 6-(4-chlorophenyl)-4,6-dioxohexanoic acid (0.49 g, 1.93 mmol) and triethylamine (0.31 ml, 2.31 mmol) in MeOH (30 ml) was stirred overnight at room temperature. The reaction was quenched with 5% HCl aqueous solution (pH2), then extracted with DCM (3×10 ml). The combined organic layers were dried over Na_2_SO_4_, filtered and concentrated. The residue was purified by column chromatography on silica gel (20 to 60% AcOEt in hexanes) to afford 3-(5-(4-chlorophenyl)-1-(4-methoxyphenyl)-1H-pyrazol-3-yl)propanoic acid (285 mg, 42% yield). LCMS: MS(ESINeg) m/z = 355 [M-H]^-. 1^H NMR (400 MHz, Chloroform-*d*) δ 10.25 (brs, 1H), 7.27 (d, *J* = 8.5 Hz, 2H), 7.20-7.12 (m, 4H), 6.87 (d, *J* = 8.9 Hz, 2H), 6.35 (s, 1H), 3.82 (s, 3H), 3.07 (t, *J* = 7.5 Hz, 2H), 2.83 (t, *J* = 7.5 Hz, 2H). Compound purity of >97% was quantified using mass spectrometry analysis.

### Synthesis of Bis(ethylmaltolato)oxovanadium(IV) (BEOV)

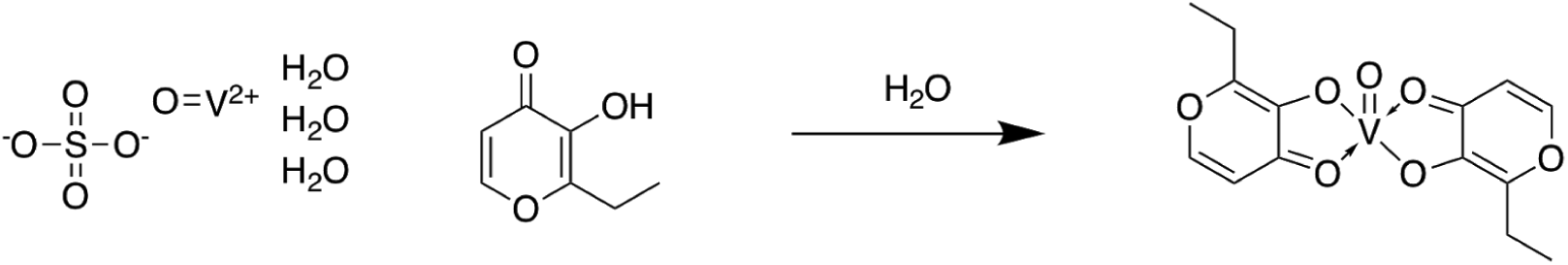

Vanadyl sulfate trihydrate (25 g, 115 mmol), dissolved in 25 mL water was added to ethylmaltol (43.4 g, 310mmol) dissolved in 125 mL hot water under argon, and the resulting solution was heated gently with stirring for 30 min. The pH was adjusted very slowly to 8.5 by addition of NaOH (12.7 g, 319 mmol) in 10 mL water. The resulting mixture was then refluxed for 2 h, and then allowed to cool to room temperature. The dark blue-grey solid was collected by vacuum filtration, washed with cold water, and dried *in vacuo* to produce bis(ethylmaltolato)oxovanadium(IV) (BEOV) with yield of 88%. The compound was analyzed by mass spectroscopy to yield a 93% purity.

### Cell lines

LS1034, HeLa, HEK293T cells were purchased from the American Type Culture Collection. REC1, SF295, OVISE, COLO320, BEN, HT29 and SNU449 were provided by the Broad-Novartis Cancer Cell Line Encyclopedia. Wildtype and MDR1 overexpressing Kuramochi cell lines were gifts from Elizabeth Stover. HeLa PDE3A CRISPR KO cells (PDE3A-/-cells) were previously described^15^. LS1034, SF295, A2058, COLO320, SNU449 and OVISE cell lines derived with Cas9 were provided by the Broad Cancer Dependency Map. LS1034, REC1, OVISE, SF295, COLO320, SNU449 and Kuramochi cell lines were cultured in RPMI (Thermo #11875093). A2058 and Hela cell lines were cultured using DMEM (Thermo #10566016). All media was supplemented with 10% heat inactivated serum FBS (Sigma 18A079) and 1% penicillin-streptomycin G (Thermo#10378016) with the exception of HEK293 which was maintained without antibiotics.

We have compared STR profiles of the cell lines in our PRISM collection with established and published STR profiles reported by vendors and in literature. Misidentified cell lines or other STR conflicts are listed in Supplementary Table 13.

For validation studies, RNAseq gene expression data was obtained from the CCLE website (https://portals.broadinstitute.org/ccle). Identity of parental human cell lines was confirmed by STR fingerprinting (Genetica) and were confirmed negative for mycoplasma using the MycoScope™ PCR Mycoplasma Detection Kit (Genlantis).

### Cloning

pXPR_003 and pXPR_023 vectors were acquired from the Broad Genetic Perturbation Platform (GPP). Oligos for sgRNAs designs were generated using Broad GPP sgRNA guide generator resource (https://www.broadinstitute.org/gpp/db/analysis-tools/sgrna-design) and the respective oligos were synthesized by Integrated DNA Technologies. Respective guide sequences can be found in the table below. To clone the sgRNAs into either the pXPR_003 guide-only or pXPR_023 all-in-one CRISPR lentiviral expression systems, we followed the protocol available on the Broad GPP website (https://www.broadinstitute.org/gpp/db/resources/protocols). CRISPR sgRNA sequences are shown in Supplementary Table 14.

### Lentiviral packaging and infection

In order to generate lentiviral vectors for CRISPR-Cas9 gene knockout, HEK293T cells were seeded in 6 well plates at a density of 1.5E6 cells per well. Cells were then transfected with a mixture of TransIT®-LT1 Transfection Reagent (MirusBio#MIR2304), psPAX2, pMD2.G, and lentiviral plasmid diluted in Opti-MEM™ (Thermo#31985062). The following day media was changed to DMEM (Thermo #10566016) with 30% FBS (Sigma 18A079). 72 hours after transfection, media containing newly produced virus was collected and run through a 0.45 µM filter to remove cellular debris.

For mammalian cell infection, a mixture of 3E6 cells were infected per single well of a 12 well plate. Per well, cells, desired concentration of lentivirus, and 4 µg/mL polybrene (Millipore) were brought to a total volume of 2 mL using cell culture medium with 10% FBS. The plates were centrifuged at 2000 RPM for 2 hours at 30C. After centrifugation, 2 mL of fresh media was added to each well and the cells were allowed to incubate at 37C overnight. The following day cells were selected with puromycin for 3-10 days, until the non-infected control cells were non-viable. For validation of CRISPR-Cas9-mediated knockout, protein targets were validated by western blot and/or indel frequency was quantified by NGS.

### Western Immunoblotting

Adherent cells were washed once with cold 1x PBS (Corning #01018002) and lysed with RIPA buffer (Sigma# R0278) supplemented by protease and phosphatase inhibitors (Sigma). Protein content was quantified using the DC Protein Assay (BioRad #5000111). Samples were boiled and reduced in a mixture containing NuPAGE® Sample Reducing Agent (10X) and 4X Protein Loading Buffer before being resolved by SDS gel electrophoresis on 4-20% Tris/glycine gels (Invitrogen). Proteins were transferred onto a nitrocellulose membrane utilizing a Mini Trans-Blot Electrophoretic Transfer Cell (Bio-Rad) in Tris-Glycine Buffer (Bio-Rad) with 10% methanol for 4 hours at 60V and 4C. The membranes were then blocked in Odyssey Blocking Buffer (LI-COR #927-40000) for one hour, and then probed overnight with primary antibodies diluted in blocking buffer. The following day, membranes were washed 3 x 5 mins with 1X TBST and then probed with IRDye secondary antibodies (LI-COR goat anti-mouse 680LT 926-68020 and goat anti-rabbit 800CW 926-32211) at 1:5000 dilution for 1 hour at room temperature. Membranes were washed an additional 3 x 5 mins in 1XTBST and then imaged using the LI-COR Odyssey CLX Infrared imager.

### Genomic DNA PCR and NGS to quantify CRISPR editing frequency

To confirm efficient CRISPR cutting at target loci, PCR primers were designed flanking the sgRNA cut site by 75-100 base pairs on either side. Human genomic DNA PCR primer sequences corresponding to the regions targeted by MDR1, SLC26A2 and MTF1 sgRNAs are shown in Supplementary Table 15. Genomic DNA was isolated from knockout cell lines using the Gentra Puregene Kit (QIAGEN #158388) and amplified with the primers to yield an amplicon roughly 150-250 base pairs in length using Herculase II Fusion Polymerase (Agilent #600675). The PCR protocol involved a 2 minute denaturation at 95C, followed by 24 cycles of 95C for 2 minutes, 55C for 2 min, and 72C for 1 min. PCR samples were purified and submitted for the NGS CRISPR sequencing assay at the Massachusetts General Hospital DNA Core. Knockout efficiency was assessed by the percentage of reads containing a frameshift caused by an indel compared to the percentage of wild-type reads. Results for each guide were averaged across primer sets with successful amplification.

### CellTiter-Glo® Cellular Viability Assay

Cell viability was assayed using a modified manufacturer’s protocol for CellTiter-Glo® (Promega #G7573). Cells were seeded at a density of 2000 cells per well in a 96-well black, clear bottom plate (Corning# 89091-012) in 100 µL total media per well. The following day, different concentrations of compounds at various doses were printed in triplicate in a random-well format using the Tecan D300e Digital Dispenser. After 72, 96, 120, or 144 hours, 60 µL of a 1:2 solution of CellTiter-Glo reagent in 1x PBS (Corning #01018002) was added per well and allowed to incubate at RT for 10 minutes. Luminescence was measured with an integration time of 0.1s using an Envision Microplate Reader (PerkinElmer #2105-0010). Biological replicates were averaged and normalized to vehicle control. Dose curves were generated using Graphpad Prism.

### PDE3A viability and rescue studies

HeLa cells (ATCC) were used for cell-killing assays and co-immunoprecipitation experiments for PDE3A and SLFN12. HeLa PDE3A CRISPR KO cells (PDE3A-/-cells) were previously described (de Waal et al., 2015). Cell-killing assays were performed as previously reported^15^. Briefly, cells were seeded into 384-well plates at 1000 cells per well and 24 hours later treated with compounds over a dose range. Cells were incubated at 37°C for 72 hours and cell viability was measured using Cell TiterGlo (Promega). Percent viability of treated wells was calculated by normalizing to the average Cell TiterGlo values from untreated wells.

### PDE3A co-immunoprecipitation assay

PDE3A immunoprecipitation and Western blotting of coprecipitated SLFN12-V5 protein experiments were performed as previously described^15^. Briefly, HeLa cells were plated onto 15 cm plates at 3×10^6^ cells per plate, and transfected the next day with 15 µg of pLX307-SLFN12 plasmid (TRC clone ID: TRCN0000476272,) using FuGene 6 at 4:1 ratio. Roughly 72 hours post-transfection, cells were treated with 10 µM of the repurposing hit compounds or 1 µM of DNMDP or 1 µM anagrelide for 6 hours. Cells were collected and lysed with a modified RIPA buffer (150 mM NaCl, 10% glycerol, 50 mM Tris-Cl pH 8.0, 50 mM MgCl_2_ 1% NP-40), supplemented with EDTA-free protease inhibitors (Roche), and PHOSTOP (Roche). Immunoprecipitation was performed using 2mg of total protein lysates and 1 µg of anti-PDE3A antibody (Bethyl Laboratory A302-740A) at 4°C overnight, followed by incubation with 7.5 µL each of Protein A- and Protein G-Dynabeads (Life Technologies 10001D and 10003D) at 4°C for 1 hour. Beads were washed with lysis buffer and bound proteins were eluted with 30 μl of LDS PAGE gel loading buffer. IP products or 60 µg input lysates were resolved on 4–12% Tris-Glycine SDS-PAGE gels and immunoblotted with an anti-V5 antibody (Life Technologies R96205, 1:5,000), Bethyl anti-PDE3A antibody (1:1,000), and secondary antibodies from LI-COR Biosciences (Cat.# 926-32210 and 926068021, each at 1:10,000). Blots were washed and imaged using a LI-COR Odyssey infrared imager.

### PDE3A and PDE3B enzyme activity assays

Recombinant PDE3A (catalog 60030) and PDE3B (catalog 60031) activity assays were performed by BPS Bioscience. The commercially available assay utilizes fluorescence polarization to assess cAMP substrate abundance following incubation with test compounds. Compounds were tested in duplicate at nine concentrations in a half-logarithmic dilution series (top concentration of 10 µM) with final DMSO concentration of 1%. The enzymatic reactions were conducted at room temperature for 60 minutes in a 50µl mixture containing PDE assay buffer, 100 nM FAM-cAMP, a PDE enzyme and the test compound. After the enzymatic reaction, 100 µl of a binding solution (1:100 dilution of the binding agent with the binding agent diluent) was added to each mix and the reaction was performed at room temperature for 15 minutes. Fluorescence intensity was measured at an excitation of 485 nm and an emission of 528 nm using a Tecan Infinite M1000 microplate reader. Fluorescence intensity was converted to fluorescence polarization using the Tecan Magellan 6 software. The fluorescence polarization data were analyzed using Graphpad Prism. The fluorescence polarization (FPt) in the absence of the compound in each data set was defined as 100% activity. In the absence of PDE and the compound, the value of fluorescent polarization (FPb) in each data set was defined as 0% activity. The percent activity in the presence of the compound was calculated according to the following equation: % activity = (FP-FPb)/(FPt-FPb)×100%, where FP= fluorescence polarization in the presence of the compound. The values of % activity versus a series of compound concentrations were then plotted using non-linear regression analysis of sigmoidal dose-response curve generated with the equation Y=B+(T-B)/1+10((LogEC50-X)×Hill Slope), where Y=percent activity, B=minimum percent activity, T=maximum percent activity, X= logarithm of compound and Hill Slope=slope factor or Hill coefficient. The IC50 value was determined by the concentration causing a half-maximal percent activity.

### Differential scanning fluorimetry (DSF/thermal shift) analysis of compound binding to PDE3A

Recombinant PDE3A (residues 677-1141) at 5 µM concentration was incubated for 20 minutes at RT in the absence and presence of 100 µM compound. The reaction buffer was 20 mM Hepes pH 7.4, 150 mM NaCl, 500 µM TCEP, and 5 mM MgCl2, and had a final concentration of 1% DMSO. After incubation, SYPRO orange (Thermo Fisher Scientific) was added to give a final concentration of 10x relative to stock concentrate. For each solution 5×10 µL aliquots were transferred to a 384 well plate and placed in a Lightcycler 480 (Roche Life Science). The temperature was increased from 25 °C to 95 °C using a gradient of 0.06 °C/second. The data was processed using the Lightcycler software. The melting temperatures and errors reported were calculated from the average of the 5 replicates.

### LS1034 CRISPR-Cas9 genome-wide knockout screen

LS1034 cells stably expressing Cas9 (pLX_311) were obtained from the Cancer Dependency Map project. The Brunello virus library, which contains 76,441 sgRNAs and 1000 control sgRNAs targeting 19,114 genes, was obtained from the Broad Genetic Perturbation Platform^45^. Before screening, Brunello virus was titered to identify the amount of virus that would yield an infection efficiency of 0.3-0.6 for 3E6 LS1034-Cas9 cells in a well of a 12-well plate.

For the screen, 432E6 LS1034-Cas9 cells were infected with Brunello virus in 12-well plates via centrifugation at 2000 rpm and 30C. The following day, cells were trypsinized and split into two biological replicates and selected with 6 µg/mL puromycin for 7 days. After selection, replicates were seeded into 16 µM tepoxalin (Wuxi) or DMSO control. Cells were maintained at 37C and 5% CO2 in CellSTACK 1272 cm^2^ 2-STACK flasks (Corning #3269) in RPMI with 10% FBS for 4 weeks. Cells were trypsinized and reseeded every 7 days at a minimum of 40 million cells per passage in order to maintain ∼500x library representation. Media and drug was refreshed every 3-4 days for a total of three weeks. Cell pellets were collected following puromycin selection and at the conclusion of the screen. Genomic DNA was isolated from cell pellets using the NucleoSpin® Blood XL Columns (Machery Nagel #740950.50) following the manufacturer’s protocol. Genomic DNA PCR and DNA sequencing was performed by the Broad Genetic Perturbation Platform and Broad Genomics Platform using standard protocols.

### LS1034 CRISPR-dCas9 genome-wide activation screen

LS1034 cells were first derivatized with pXPR_109 using lentiviral transduction in order to stably express dCas9-VP64^46^. The Calabrese B virus library, containing 56,476 sgRNAs and 500 controls targeting 18,843 genes, was obtained from the Broad Genetic Perturbation Platform. CRISPR guide RNAs were delivered using pXPR_502, which encodes PP7-P65-HSF and an sgRNA. To confirm dCas9-VP64 activity, LS1034-dCas9-VP64 were infected with control sgRNAs targeting CD4 (BRDN0002435409; guide sequence: ATGTTCCCTGAGAGCCTGGG) or CD45 (BRDN0002435411; guide sequence: GTTGTTCTAAGTCAGTAGAA). Following puromycin selection, cells were stained with Brilliant Violet 421 anti-CD45 (Biolegend 368522) and APC anti-human CD4 (Biolegend 300514). 68% of sgCD45-transduced cells expressed CD45 and 28% of sgCD4-transduced cells expressed CD4.

For the screen, 288E6 cells were infected with the Calabrese B virus library via centrifugation as described above. Average observed infection efficiency was 0.31. The following day, cells were split into two replicates and re-seeded in CellSTACK 1272 cm^2^ 2-STACK flasks (Corning #3269). After allowing cells to adhere overnight, media was changed to introduce 6 µg/mL of puromycin selection for 6 days. After selection, replicates were split into DMSO or 16 µM tepoxalin (Wuxi) drug arms in duplicate, and cultured for two weeks as described above. Genomic DNA was harvested and sequenced as described above.

Brunello and Calabrese CRISPR guide virus libraries were obtained from the Broad Genomic Perturbation Platform (also available from Addgene). The pXPR_003, pXPR_023, pXPR_109, pXPR_502, and pXPR_311 vectors were gifts of John Doench.

### CRISPR Screen Analysis

For analysis of the CRISPR and CRISPRa data, we first filtered out the guides from the sgRNA library that are known to target more than a single gene or had counts of less than 50 in the initial pDNA pool. Then, the cell counts for each guide under treatment are normalized with respect to total library size. The logarithms of the ratio between the normalized counts in the treatment compared to the control (DMSO) condition for each guide were computed to yield guide-level log fold-changes in cell counts. Gene-level log fold-changes were computed using the average of the log fold-changes of all the guides targeting each gene. Next, the treatment and control conditions were compared using the MAGeCK-MLE method, using the suggested number of permutation rounds (10), to compute the statistical significance of the gene level log fold-change of the normalized cell counts in the treatment with respect to DMSO^47^. These log fold-changes were plotted against the false discovery rates resulting from the Benjamini-Hochberg correction of the two-sided p-values produced by the MAGeCK-MLE calculation (Fig. 4).

### Tepoxalin Competition Assay

3E6 LS1034 Cas9 cells stably expressing Firefly luciferase and guides to *ABCB1* or an intergenic region on chromosome 2 (cutting control guide) were co-cultured with 3E6 LS1034 cells stabling expressing Renilla luciferase. Cells were cultured in RPMI containing 10% FBS in a 10 cm tissue culture plate and treated with either 16 µM tepoxalin (Wuxi) or vehicle control. The co-culture was passaged every 4 days and re-seeded in media with respective treatment at a bottleneck of 6E6 cells. Luciferase activity was quantified in triplicate using Dual-Glo® Luciferase Assay System (Promega #E2920) and measured using an Envision plate reader (Perkin-Elmer) with ultrasensitive luminescence. Data points were collected at day 0 and every 4 days until 12 days of treatment. The Firefly to Renilla luminescence ratio was calculated and normalized to the initial Day 0 measurement. The results are representative of three independent experiments.

### Tepoxalin cell permeability assay

Compound was incubated in naïve cell media (no cell control, or NCC) or in the presence of cells at 1 million cells/mL. The incubation was carried out in a 37°C with 5% CO2 cell incubator with gentle plate shaking for 3 hours. Following the incubation, the NCC samples were placed on the bench until final analysis. Cell samples were centrifuged at 500 x g for 5 minutes. The supernatant was removed, and the cells were washed twice with cold PBS. Cells were again centrifuged at 500 x g for 5 minutes, the supernatant was removed and discarded, and the cells were resuspended in 130 µL water (Honeywell, catalog #33015-1L). The entire volume was then transferred into a Covaris microTUBE plate (Covaris, catalog #220078) and sonicated on a Covaris LE220 focused-ultrasonicator. Samples were transferred to a 96-well plate and centrifuged at 3000 x g for 15 min at 20°C. The supernatant was transferred to a new 96-well plate. For both the NCC and cell samples, 5 µL was transferred to a 96-well plate. To each sample was added 45 µL cell media, 50 µL water, and 50 µL acetonitrile containing internal standard. Samples were centrifuged again, and a final 100 µL aliquot was transferred to a 96-well plate for analysis.

Samples were analyzed on a UPLC-MS/MS system consisting of a Waters Acquity I-Class FTN and AB Sciex 4500 Triple Quad mass spectrometer with compounds detected by positive mode MRM detection. Mobile phase A consisted of water with 0.1% formic acid (Honeywell, catalog #33015-1L), while mobile phase B consisted of acetonitrile with 0.1% formic acid. The gradient ran from 10-95% B over 0.8 minutes at a flow rate of 0.9 mL/minute. An Acquity BEH C18, 1.7 m, 2.1 x 50 mm column (Waters, P/N 186002350) was used with column temperature maintained at 65°C. Sample concentrations were determined using a standard curve and dilution quality-control samples prepared in a surrogate matrix. Analyst 1.6.2 software was used for integration and calculation determination.

### Tepoxalin stability assay

1 µM of tepoxalin was added to PBS, RPMI with 10% FBS, or acetonitrile in duplicate and measured by mass spectrometry with time points prepared at 0, 24, 48, and 72h. The mass spectrometer was run in positive mode using MRM detection for tepoxalin and the internal standard (75 nM midazolam).

### Tepoxalin drug synergy/antagonism assays

Tepoxalin or paclitaxel were added in a dose response matrix format to LS1034 or REC1 cells, and viability was assessed by CellTiter-Glo. For drug combination viability data, we first normalized CellTiter-Glo measurements by the median over DMSO wells on each plate. We then used the R package synergyfinder ^48^ to estimate Bliss synergy scores across all dose combinations, applying synergyfinder’s default baseline correction method. Synergy values for each drug combination and cell line were summarized by the synergy score with highest magnitude across dose combinations (max synergy). We verified that qualitatively similar results were obtained using other synergy models (i.e. Loewe, HSA, ZIP), and methods for aggregating synergy scores across dose combinations, such as averaging.

### P-glycoprotein transporter cell based antagonism assay (Calcein AM)

The P-glycoprotein antagonism assay for tepoxalin and RWJ20142 was performed by Eurofins using their standard protocols. In brief, MDR1-MDCK cells are seeded in a 96-well culture plate at 30,000 cells/well and are used on days 2 or 3 post-seeding. On the day of assay, the test compound is prepared in assay buffer (HBSS-HEPES, pH 7.4), added to the cell plate, and pre-incubated at 37 C for 15 min. Subsequently, calcein AM is added to the plate followed by a 20-minute incubation at 37 C. The plate is then washed with cold assay buffer followed by fluorescence reading. The percent of control activity is calculated by comparing the signal in the presence of the test compound to the vehicle control. Subsequently, percent inhibition is calculated by subtracting the percent activity by 100.

### MDR1-MDCK permeability assay

The ability of tepoxalin to inhibit loperamide transport across a monolayer was assessed by Cyprotex in the MDR1-MDCK permeability assay following standard protocols. In brief, MDR1-MDCK cells are cultured for a 4-day period to form a monolayer. Once a monolayer is formed, loperamide, a known MDR1 substrate, is added to the apical side of the membrane and transport to the basal side of the membrane is quantified over a 60-minute time period using mass spectrometry detection. Inhibition of MDR1-mediated transport of loperamide across the monolayer was then assessed by adding a dose range of tepoxalin or positive control (verapamil). The percent of control activity is calculated by comparing the baseline MDR1 loperamide transport with drug-treated groups. Subsequently, percent inhibition is calculated by subtracting the percent activity from 100.

### Transcriptional profiling by RNAseq

8E5 cells LS1034 cells were seeded in 12-well plates (VWR International #29442-040) and incubated at 37C overnight. The following day, cells were treated in triplicate with 12 µM tepoxalin or DMSO vehicle control for 6 hours. All cell pellets were collected at the same time via trypsinization followed by centrifugation at 300xg for 5 minutes and washed with 1X PBS (Corning #01018002). Immediately after pelleting, cells were isolated using the RNeasy Mini Kit (Qiagen #74104) following the manufacturer’s protocol with DNase treatment.

Wild-type SF295 cells and SF295 containing Cas9 and sgRNAs targeting GFP or MTF1 were plated in triplicate on a 6 well plate. The following day, cells were isolated using RNA RNeasy Miniprep Kit (Qiagen #74104) following the manufacturer’s protocol with DNase treatment. RNA quality was confirmed by Bioanalyzer (Agilent). Library preparation was performed by the Dana-Farber Molecular Biology Core Facility using the KAPA mRNA HyperPrep Kit (Roche). Nucleic acid was sequenced using an Illumina NextSeq 500 instrument (PE75 kit).

Gene level expression values were obtained from RNA sequencing using the TOPMed RNA-seq pipeline (https://github.com/broadinstitute/gtex-pipeline/blob/master/TOPMed_RNAseq_pipeline.md). In brief, FASTQs are aligned to the transcriptome using the STAR alignment algorithm. RSEM was used to generate transcripts per million (TPM) gene-level expression quantifications. These tools were run using the FireCloud and Terra platform^49^. The differential expression analysis between the treatment and control (DMSO) conditions is performed using the DESeq2 R package^50^. The sequencing data is available from the Gene Expression Omnibus (GEO, accession number GSE133299).

---

**Extended Data Fig. 1.**
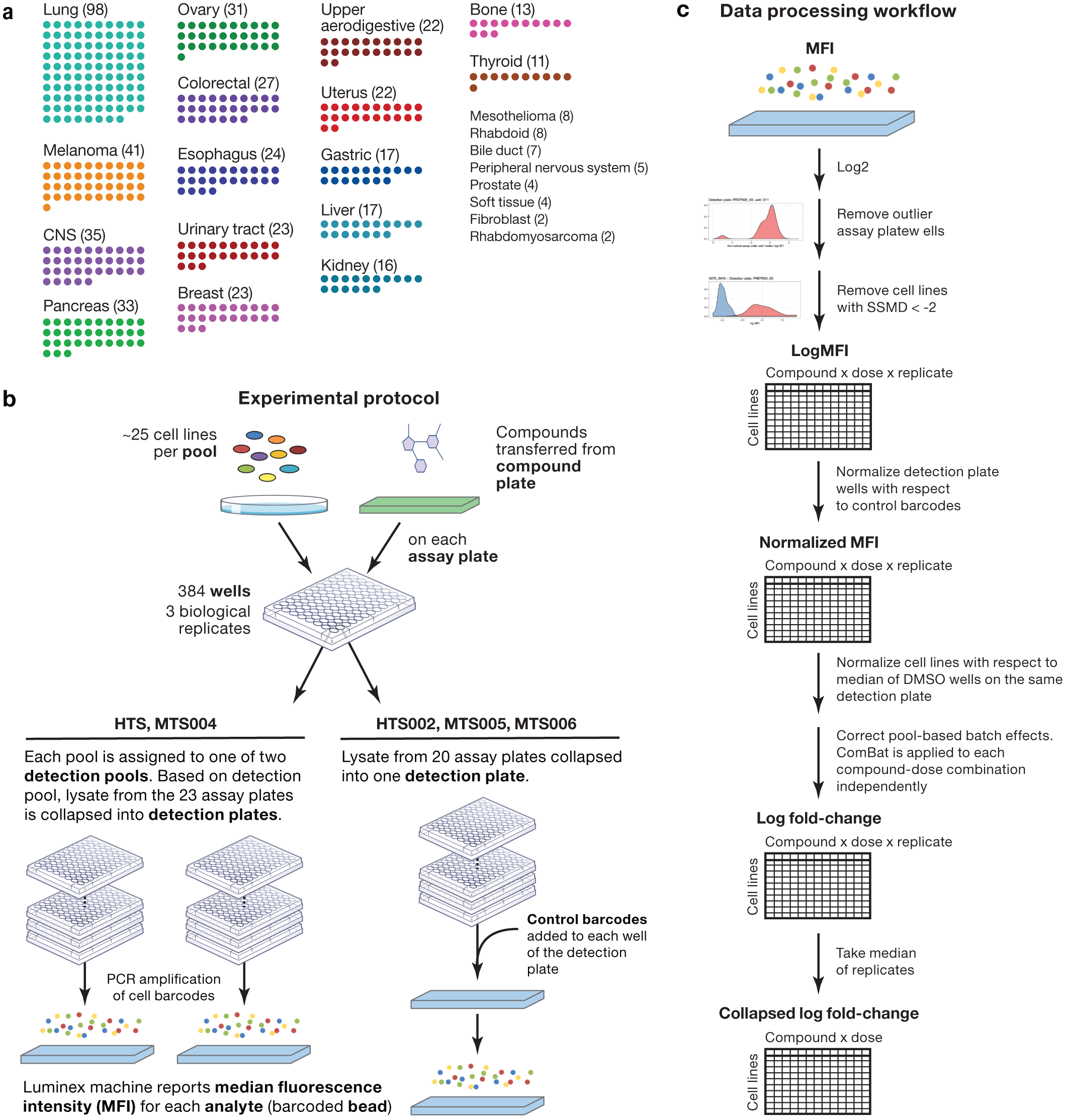
PRISM Repurposing assay and data processing overview. **a**, Lineage diversity of PRISM cell lines. The 489+ cancer cell lines tested span more than 23 tumor types. Lineages with fewer than 10 cell lines are listed on the right. **b**, Experimental protocol. Cell lines are grouped by doubling time into pools of approximately 25 cell lines. One pool is plated onto each assay plate. Compounds are transferred by pin transfer from a source compound plate (HTS and HTS002 screens) or cells are plated directly onto assay-ready plates generated by acoustic dispensing of compounds (MTS004, MTS005, and MTS006 screens). In either case, compound plates are shared by all replicates of each treatment condition. After incubation and lysis, all assay plates generated by a given compound plate are grouped and collapsed into three (HTS002, MTS005, and MTS006 screens) or six detection plates (HTS, MTS004 screens) so that each detection plate receives one or zero copies of each pool. Ten control barcodes are then spiked into each detection plate well (HTS002, MTS005, and MTS006 screens). Detection plates are amplified by PCR and detected using Luminex FLEXMAP 3D instruments. **c**, Data processing workflow. Median Fluorescence Intensity (MFI) values are calculated from fluorescence values for each replicate-condition-cell line combination and are log2-transformed. Assay plates wells are normalized, median collapsed, and compared to the normalized medians of other assay plate wells in the same well position that have been dosed by the same compound plate. A robust z-score is calculated and assay plate wells with a |z-score| > 5 are filtered. Strictly standardized mean differences (SSMD) are calculated between positive and negative control conditions for each cell line on each assay plate. Cell line-assay plate combinations with SSMD < 2 are filtered by a control-separation filter to generate the log MFI data matrix. In datasets with control barcodes added, data are normalized with respect to the median of control barcodes to generate the MFI normalized data matrix. Data are DMSO-normalized and pooling artifacts are corrected using ComBat to generate the log fold-change data matrix. Replicates are median collapsed to generate the collapsed log fold-change data matrix.

**Extended Data Fig. 2.**
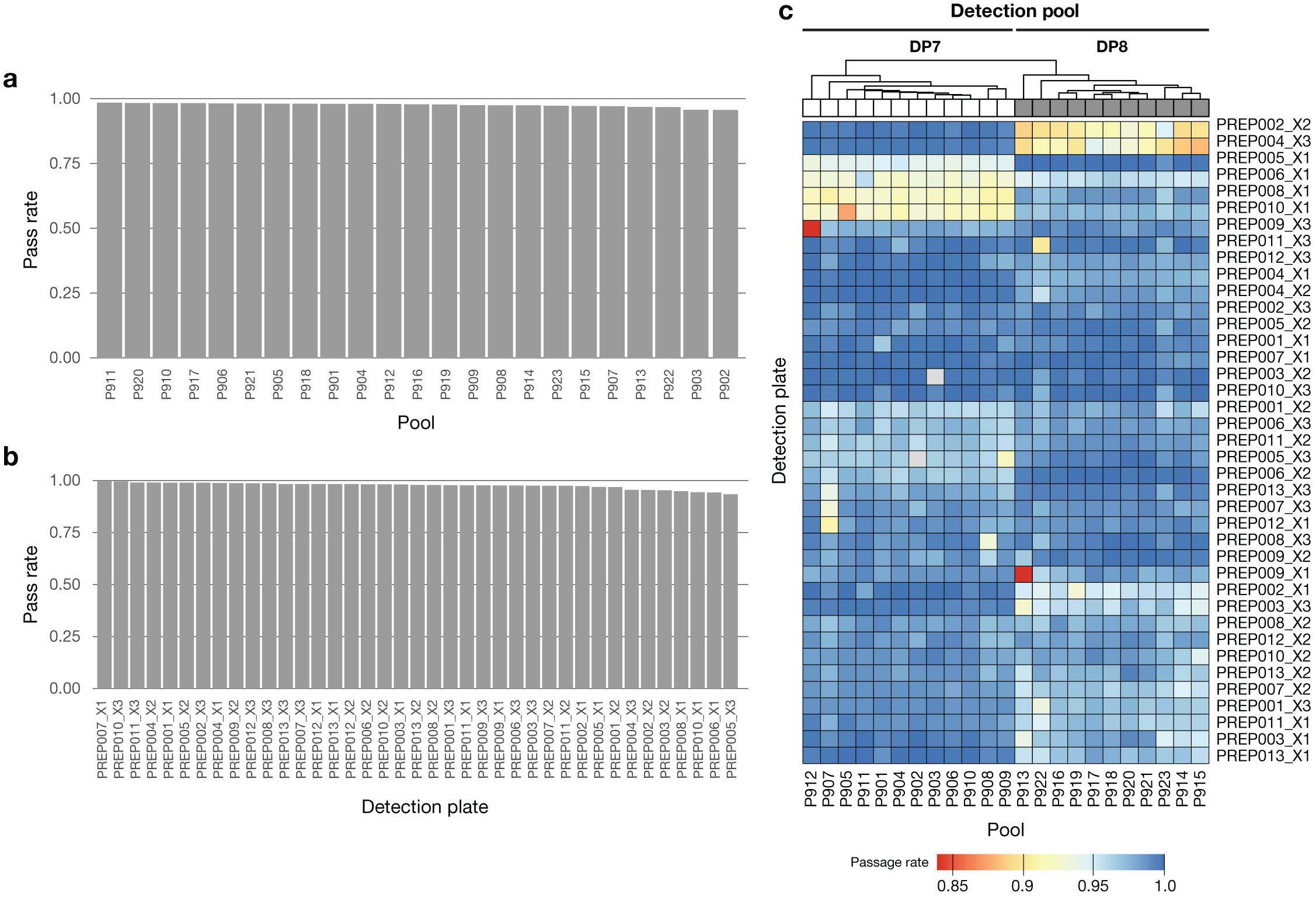
Outlier pool QC filter applied to the PRISM primary screen to detect pool-level failures. **a**, QC pass rate by pool. The fraction of treated assay plate-wells that pass the outlier filter is indicated. Cell line log MFI data are median-centered and the medians of assay plate-wells are compared within each well-detection plate combination. Extreme outliers with |robust z-score| > 5 are filtered. **b**, QC pass rate by detection plate. **c**, QC pass rate of assay plates. Overall pass rate was high (median 98.6%, minimum 83.8%). Some detection plates show higher failure rates for specific detection pools, implying failure at final detection step.

**Extended Data Fig. 3.**
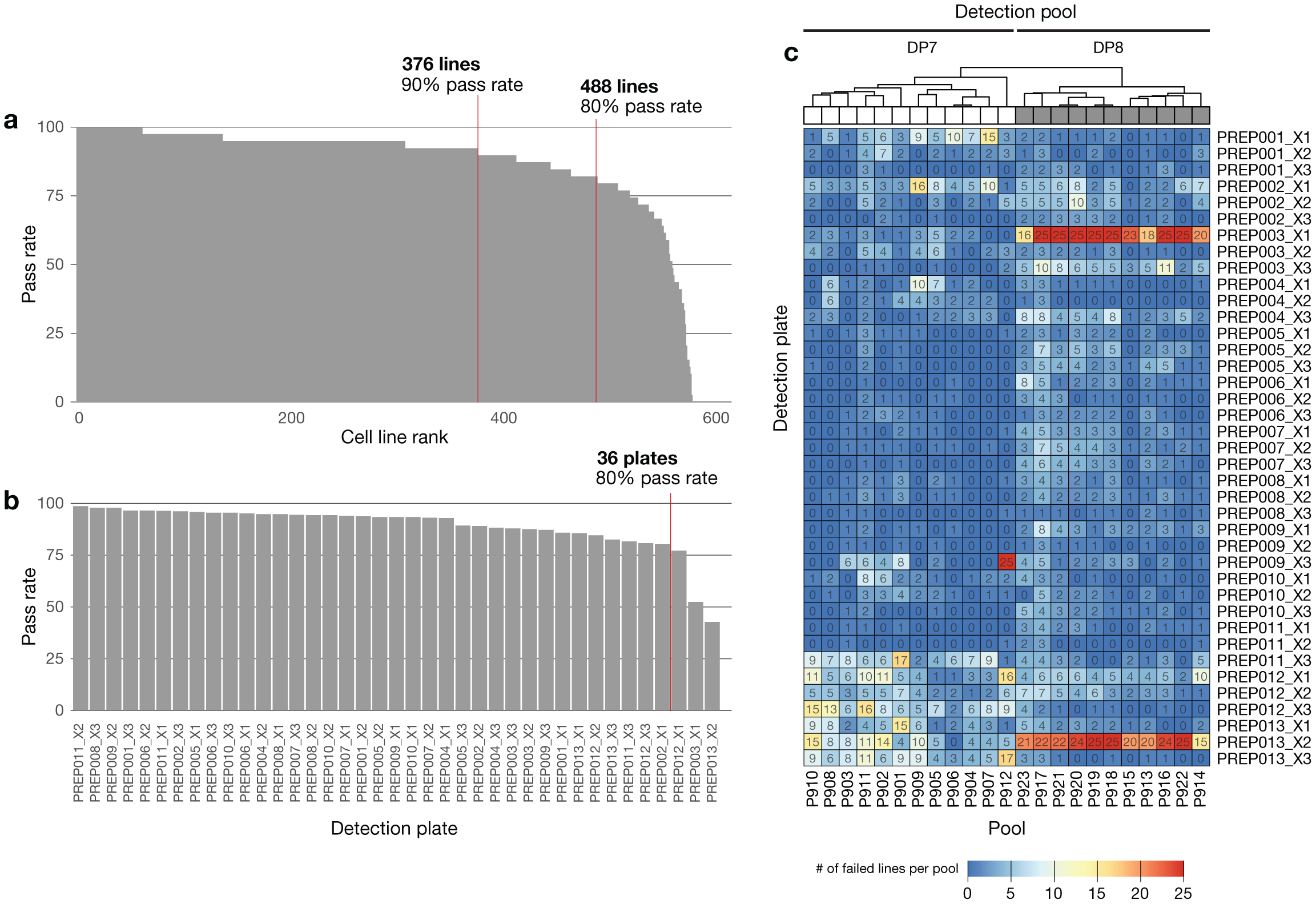
Control-separation QC filter applied to the PRISM primary screen to detect cell line failures. **a**, QC pass rate by cell line. The Strictly Standardized Mean Difference (SSMD) of log MFI values is calculated between positive control and negative control treatments for each cell line on each plate. Data from cell line-plates with SSMD < 2 are filtered. **b**, QC pass rate by detection plate. **c**, QC pass rate of assay plates. Overall pass rate was high (median 99.2%, mean 94.3%). Some detection plates show higher failure rates for specific detection pools, implying failure at final detection step. The bulk of the filtered data come from two detection plate-detection pools (PREP013_X2 and PREP003_X1 in Detection Pool 8).

**Extended Data Fig. 4.**
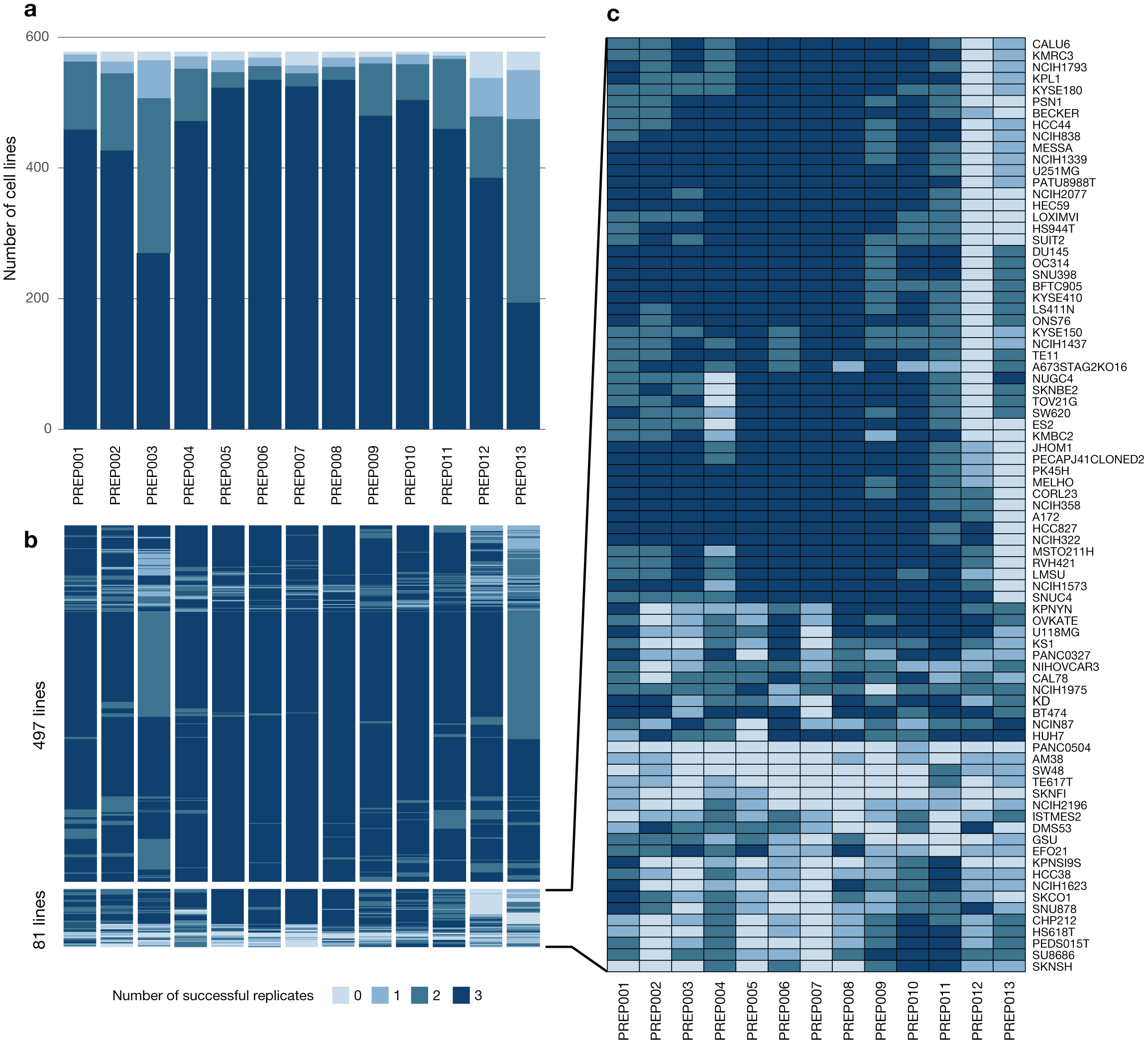
Post QC, 86% of cell lines have at least one replicate for all compounds in the PRISM primary screen. **a**, Number of technical replicates that pass the SSMD filter in the primary HTS screen, grouped by compound plate. **b**, Number of technical replicates that pass the SSMD filter in the primary HTS screen, grouped by compound plate and cell line quality. 497 cell lines have at least 1 passing replicate on each detection plate. 81 cell lines fail all three replicates on at least one detection plate. **c**, Low quality cell lines are zoomed at right, 4% of the cell lines fails across the assay, while 8% of cell lines fail on only two plates (PREP012 & PREP013).

**Extended Data Fig. 5.**
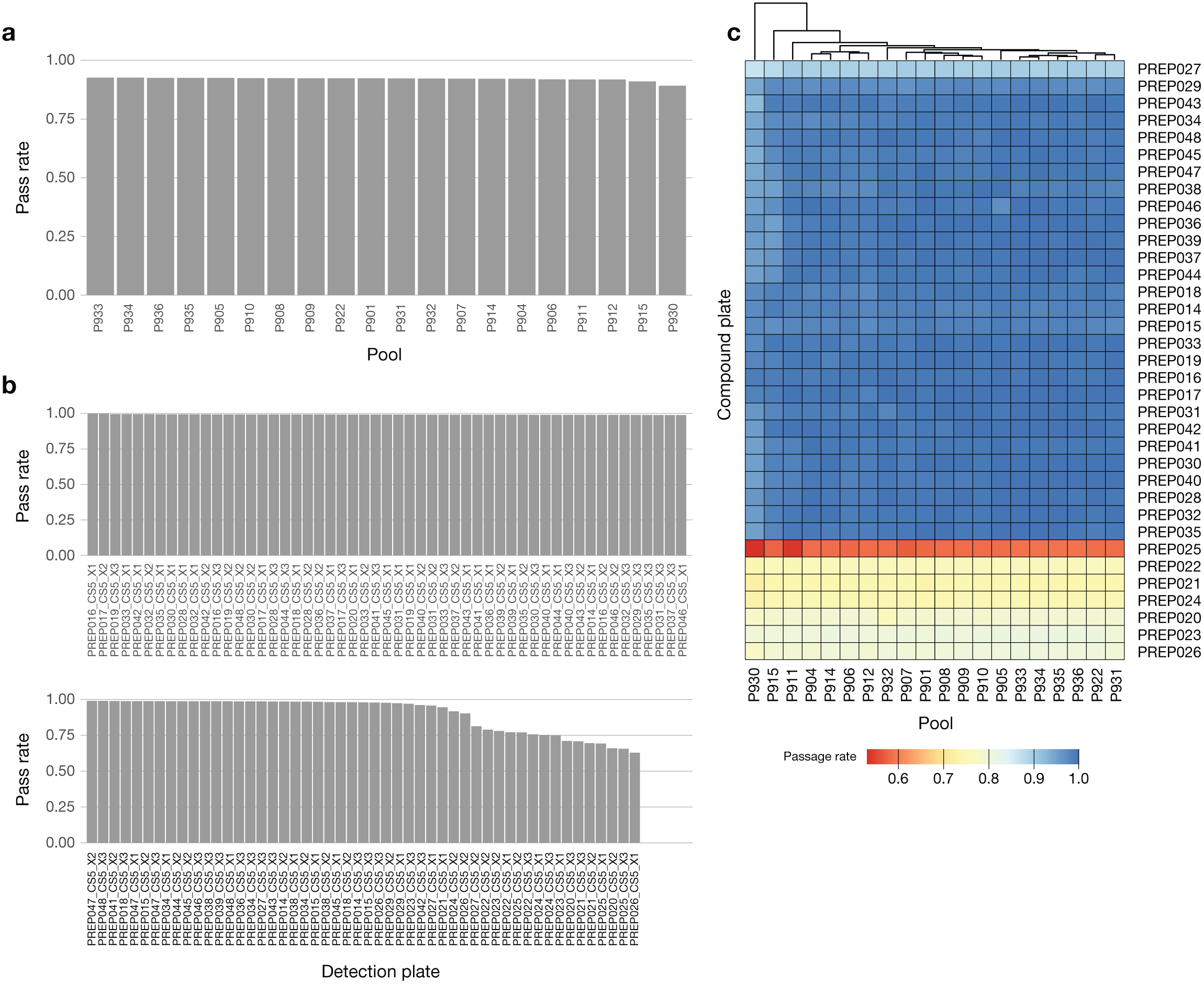
Outlier pool QC filter applied to the PRISM secondary screen to detect pool level failures. **a**, QC pass rate by pool. The fraction of treated wells that pass outlier filter is indicated. Cell line MFI data are median-centered and the pool median is compared across plates. Extreme outliers with |z-score| > 5 are filtered. **b**, QC pass rate by detection plate. The pass rate is above 95% for 81% of detection plates. **c**, QC pass rate by plate-pool combinations. Replicate plates are combined for visualization. Overall pass rate was high (median 99.1 %, mean 94.3 %), where failures are almost exclusively coming from 7 detection plates, implying failure at final detection step. Across the screen, 5.7% of the pools are filtered out as outliers.

**Extended Data Fig. 6.**
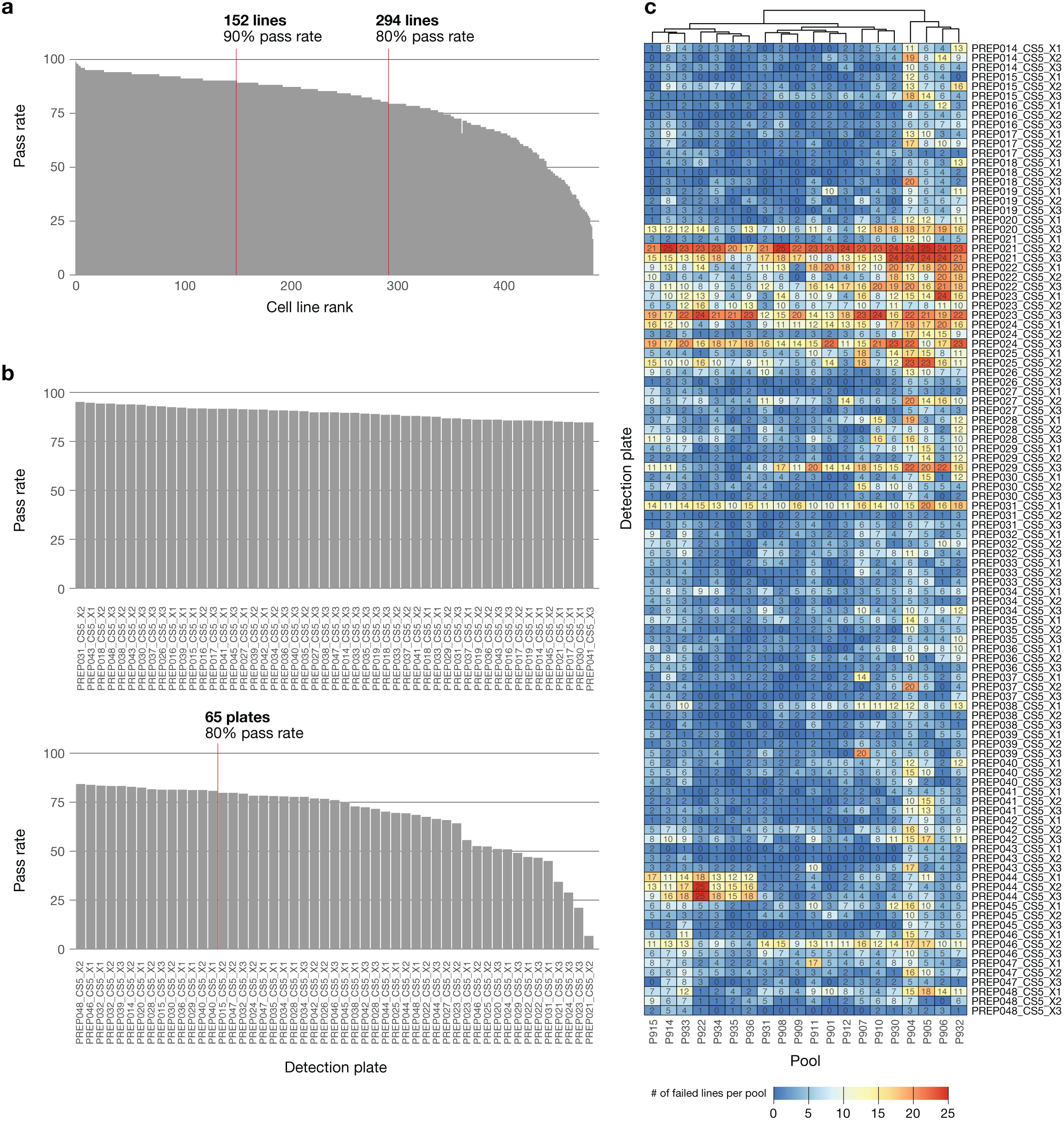
Control-separation QC filter applied to the PRISM secondary screen to detect cell line failures. **a**, QC pass rate by cell line. The Strictly Standardized Mean Difference (SSMD) of log MFI values is calculated between positive control and negative control treatments for each cell line on each assay plate. Data from cell line-plates with SSMD < 2 are filtered. **b**, QC pass rate by detection plate. **c**, QC pass rate of assay plates. Overall pass rate was lower than primary (median 99.88 %, mean 79.1 %). Similar to the primary screen, the main mode of failure is platewise failures.

**Extended Data Fig. 7.**
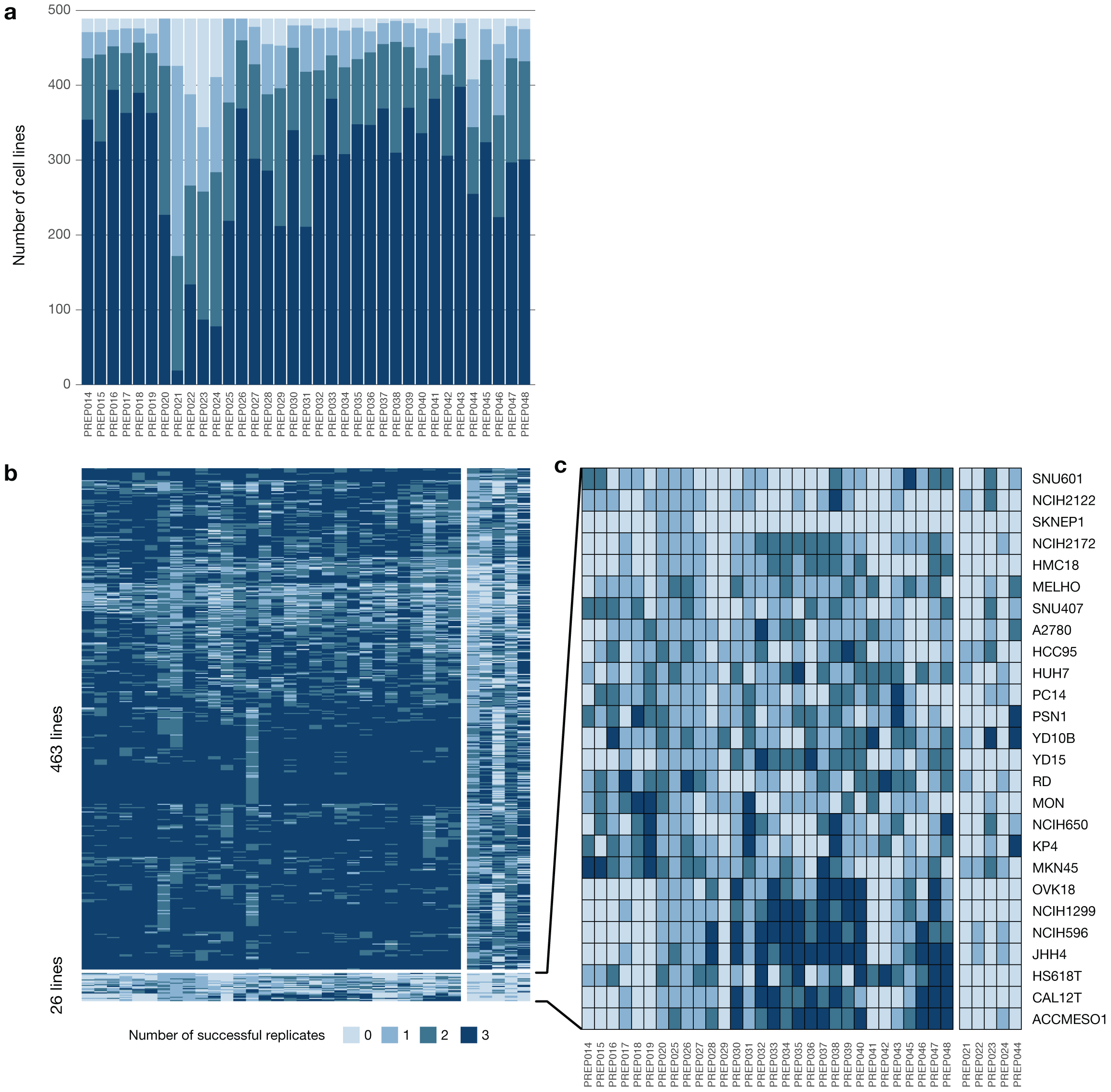
Post QC, 92% of cell lines have at least one replicate in all but five compound plates in PRISM secondary screen. **a**, Number of technical replicates that pass the SSMD filter in the secondary HTS screen, grouped by compound plate. **b**, Number of technical replicates that pass the SSMD filter in the secondary HTS screen, grouped by compound plate and cell line quality. 463 cell lines have at least 1 passing replicate on at least 85% of the compound plates. 26 cell lines fail all three replicates on at least five compound plates. **c**, Low quality cell-lines are zoomed at right, the 37% of the missing data points are coming from these 26 cell lines and 52% of them are from the worst five plates (PREP 021/022/023/024/044) (10% are both from the worst 26 cell lines and 5 plates).

**Extended Data Fig. 8.**
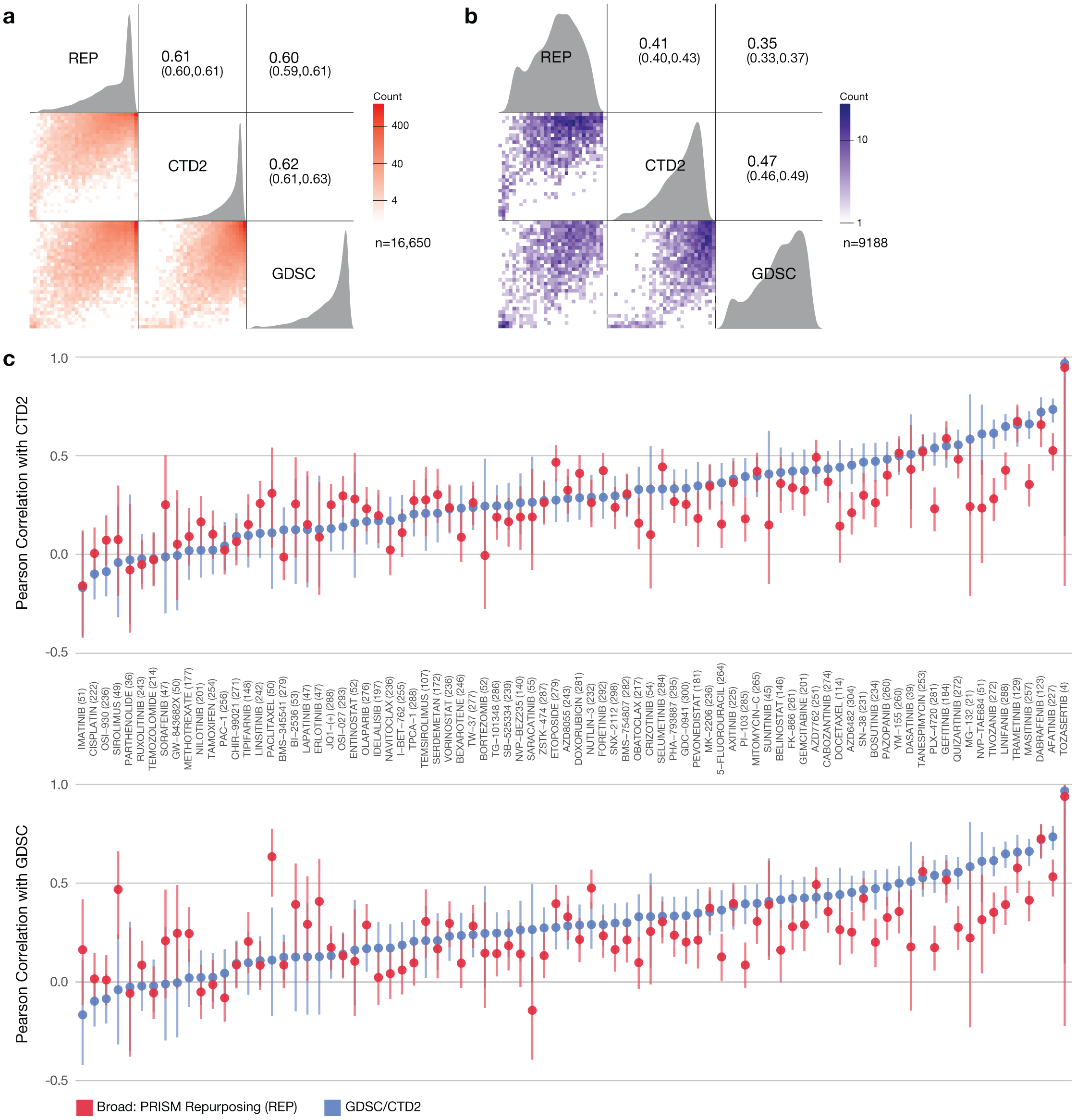
Large scale drug sensitivity datasets show modest but consistent concordance. **a**, Pairwise Pearson correlations between drug response AUCs of publicly available datasets. AUC values were recomputed for 84 compounds and 318 cell lines (median 236 cell lines per compound) over the same dose-range for all three datasets (GDSC, CTD2, REP), and capped at 1. The Pearson correlation across shared cell line-compound pairs of the datasets is above 0.6. 44.8% of cell line-compound pairs show inactivity (AUC > 0.8) in all three datasets. **b**, Pairwise Pearson correlations between drug response AUCs after removing inactive cell line-compound pairs. Pearson correlations were re-calculated after filtering out the inactive data points (AUC > 0.8 in at all three datasets). **c**, Compound-wise correlation between publicly available datasets. Correlation between PRISM data and datasets is similar to correlation between other datasets. Killing profiles are correlated across cell lines for each compound among three datasets. The Pearson correlation across datasets are given with their 95% confidence intervals, GDSC vs CTD2 (Blue) and REP vs GDSC/CTD2 (Red). Number of cell lines commonly available in all three datasets are given in parentheses after drug names. Note that the confidence intervals are particularly large for the compounds with relatively small number of cell lines. Paired t-tests on the compound-wise correlations show statistically significant (two-sided p-value: 0.012 and 0.014 for top and bottom, respectively) but small mean of differences (0.049 and 0.039 for top and bottom, respectively).

**Extended Data Fig. 9.**
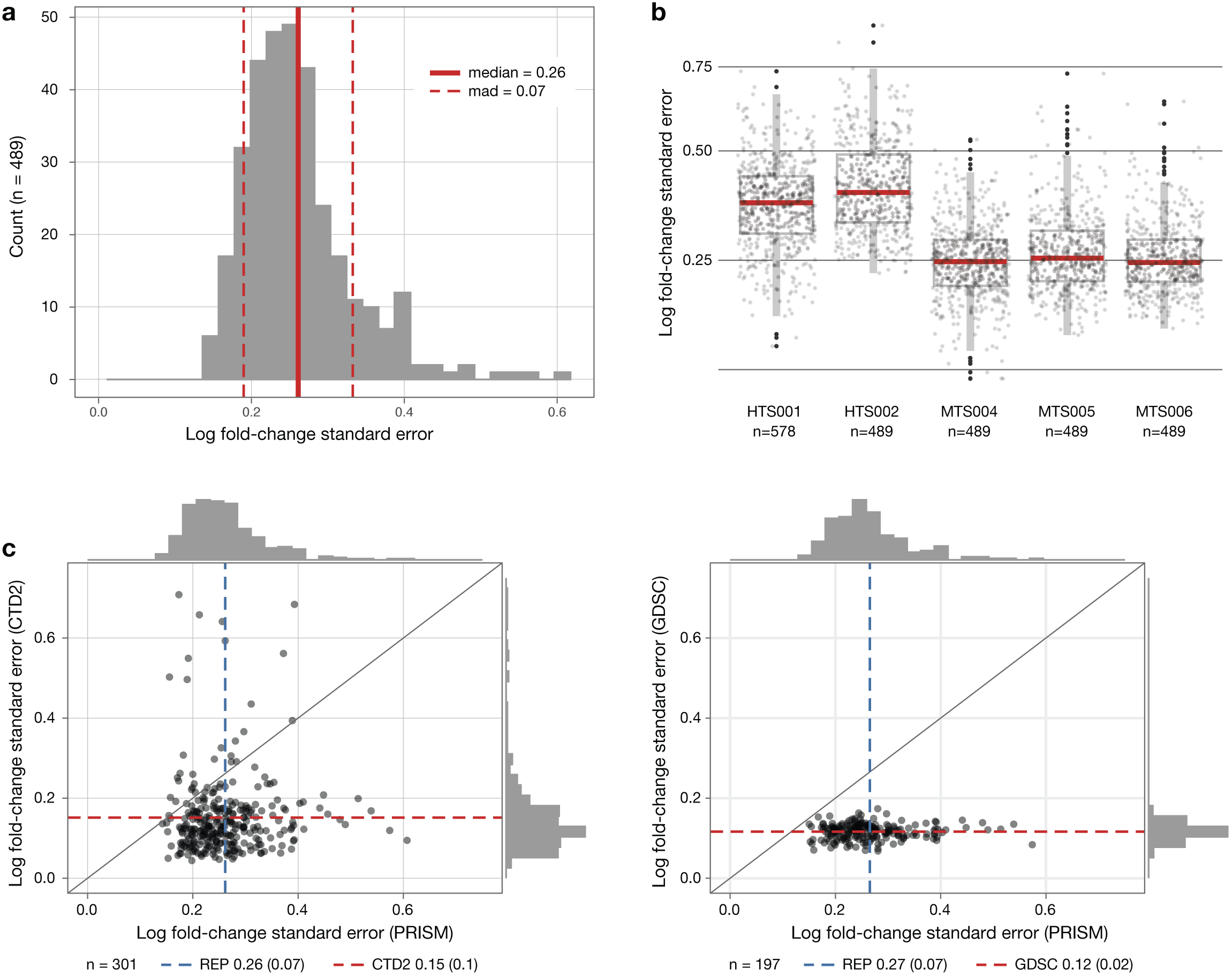
PRISM Repurposing noise quantification. **a**, Cell line standard-error estimates across vehicle-treated wells on PRISM plates treated with DMSO. Log fold-change standard errors are estimated for each cell line using DMSO-only plates included in the MTS006 screen. **b**, Comparison of standard-error estimates across screens. The error estimate calculation is repeated for each screen using DMSO wells on standard compound plates. Higher noise levels are observed in the initial high-throughput screens (HTS001 and HTS002) compared with more recently performed medium-throughput screens (MTS004, MTS005, and MTS006). **c**, Comparison of estimated standard error of vehicle control wells between high-throughput pharmacogenomic datasets. Dashed lines mark the mean standard error for each screen with the standard errors are given in parentheses.

**Extended Data Fig. 10.**
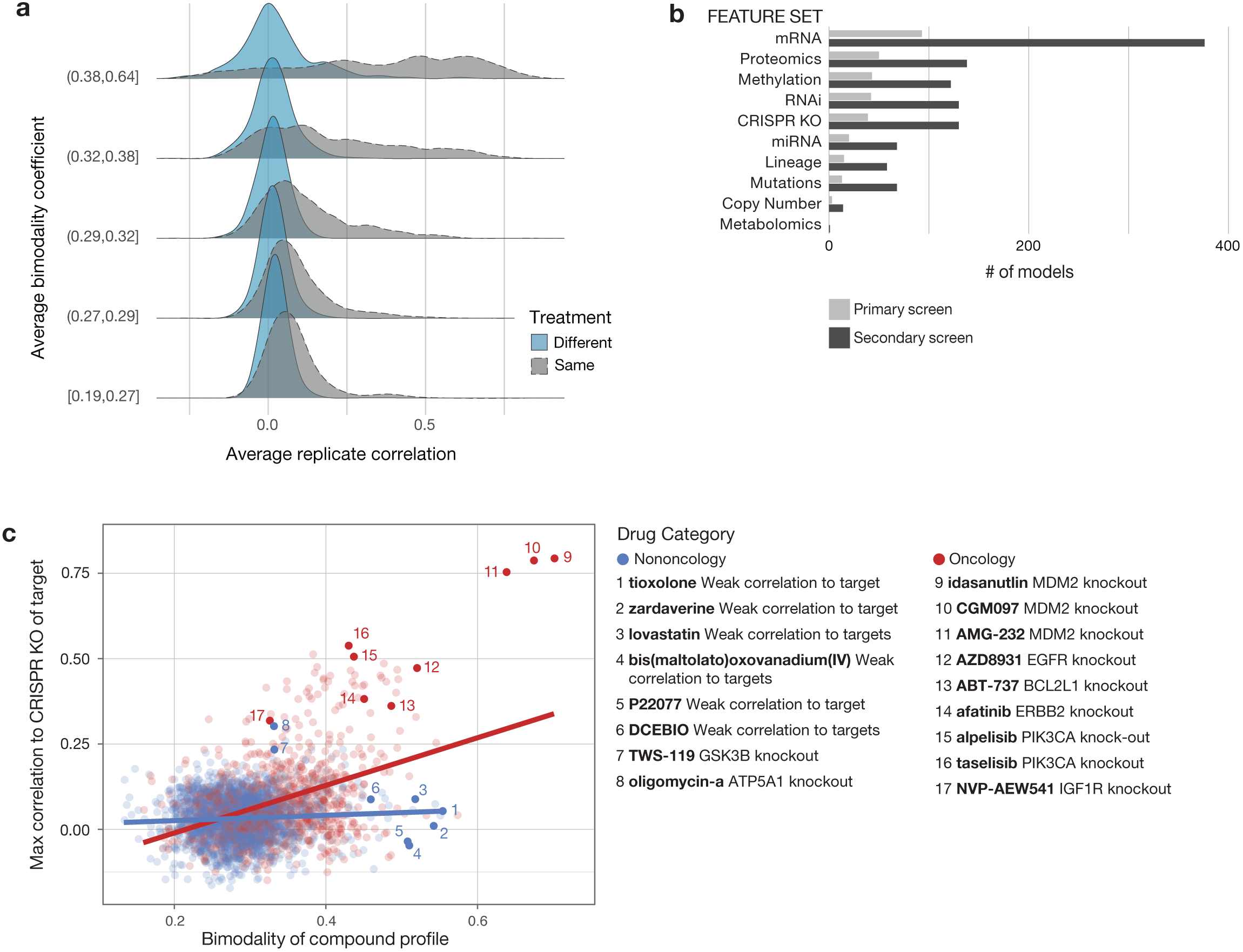
PRISM replicate correlation and comparison with genomic feature sets. **a**, Relationship between strong selective activity and replicate correlation in PRISM. The average of pairwise pearson correlations among the three replicates of each compound from the primary screen (gray) after stratifying the compounds into five equal sized sets based on the average bimodality coefficients of each profile used in this computation. For reference, the same quantities computed for randomly paired compounds and their distribution is overlaid as a null distribution (blue). **b**, Predictors of drug sensitivity based on genomic feature type. A weighted set of genomic features predictive of drug activity is determined using the ATLANTIS implementation of random forest. Baseline mRNA and protein expression (RPPA) yield largest numbers of strong predictive models in both primary and secondary screens. Models with a Pearson score greater than 0.4 are included. **c**, Relationship between bimodality coefficient of PRISM profile and correlation with CRISPR knockout of annotated gene target. Pearson correlations are computed between the primary screen PRISM log fold-change of each compound and the DepMap Avana CRISPR/Cas9 knockout gene effect scores of its known targets. The maximum correlation among all annotated targets is given as the y-axis. Bimodality coefficients are recalculated over cell lines shared by each compound and the CERES scores for its max-correlating target. Linear regression lines are shown.

**Extended Data Fig. 11.**
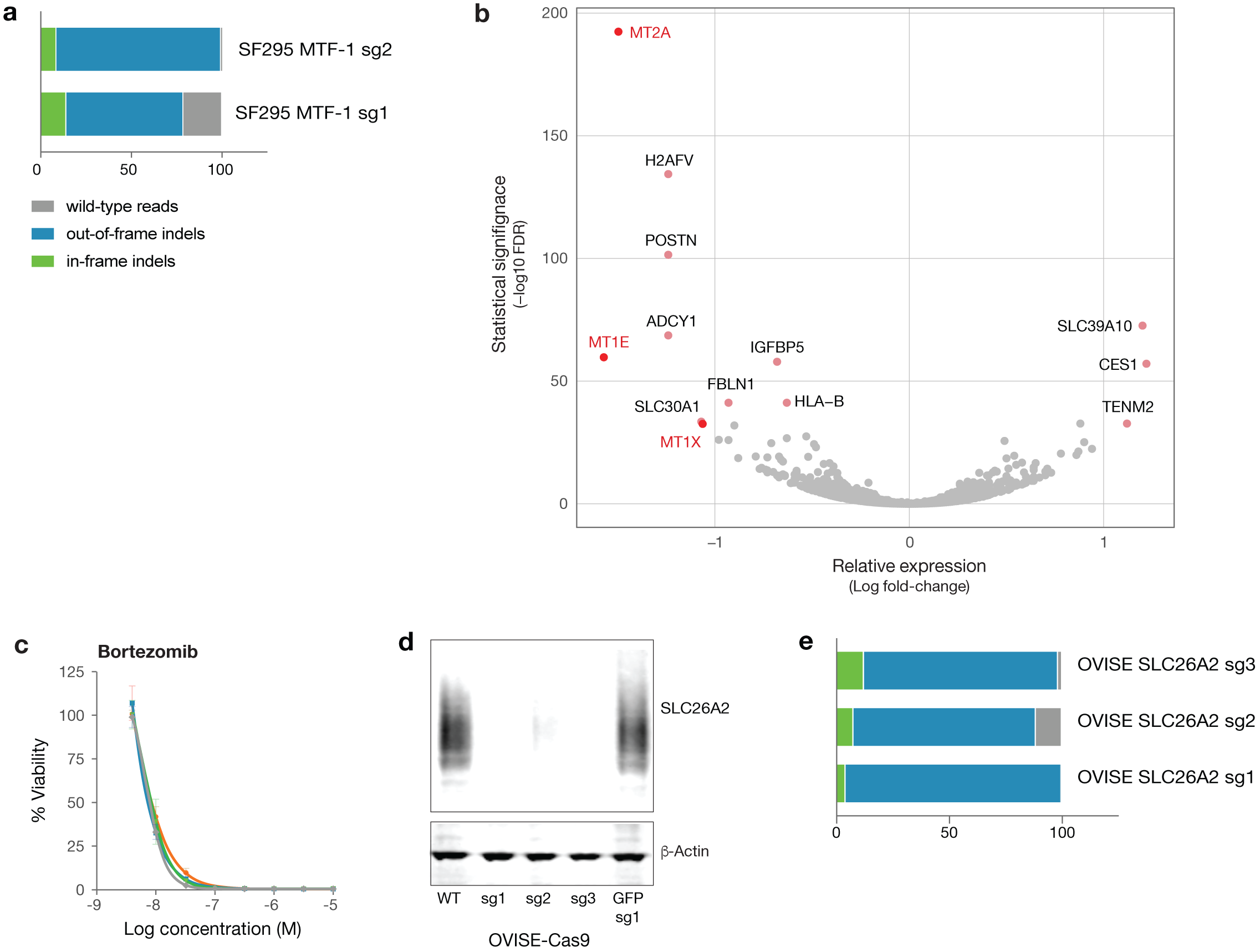
Generation of MTF-1 and SLC26A2 knockout cell lines. **a**, SF295 cells were transduced with multiple guides targeting the MTF-1 gene. Following selection, genomic DNA was isolated and the targeted region was amplified by PCR. Results from the NGS CRISPR assay are shown as percent indel formation. **b**, Differentially expressed genes in SF295 glioma cells following MTF-1 knockout by CRISPR/Cas9. Loss of MT1E, MT1X, and MT2A expression was observed upon MTF-1 knockout. Gene expression was measured by mRNA sequencing and differential gene expression was calculated by DESeq2. **c**, Drug sensitivity of SF295 cells with and without MTF-1 knockout. MTF-1 does not alter sensitivity to control chemotherapeutic bortezomib. Standard deviation across three replicates is shown with error bars. **d**, Western immunoblot validation of SLC26A2 knockout in OVISE ovarian and A2058 melanoma cancer cell lines. The SLC26A2 protein is known to migrate across a range of molecular weights due to glycosylation. **e**, OVISE cells were transduced with multiple guides targeting the SLC26A2 gene. Indel frequency at the SLC26A2 CRISPR Cas9 cut sites was assessed by NGS CRISPR assay.

**Extended Data Fig. 12.**
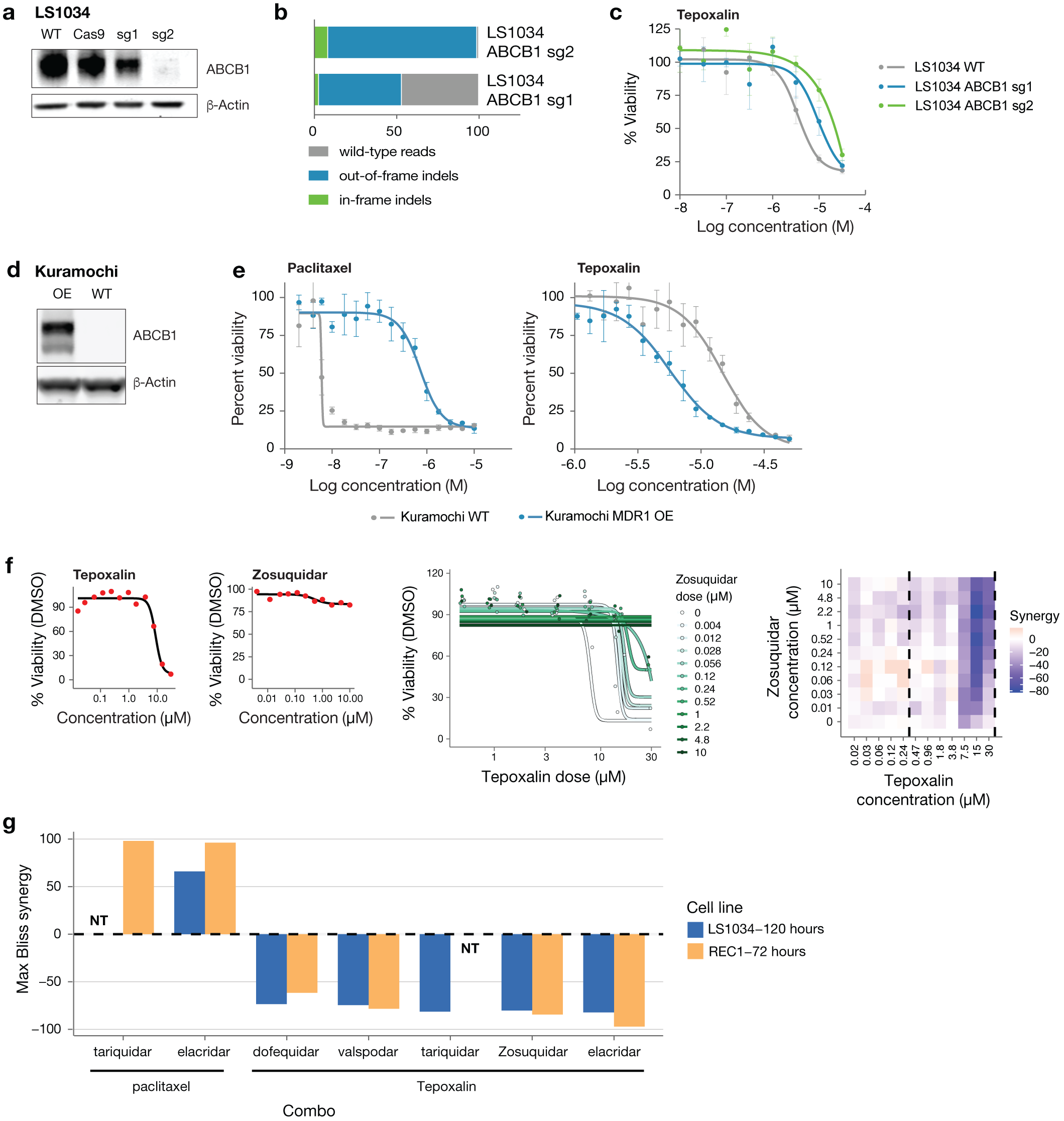
Genetic and small molecule perturbation of ABCB1 modulates response to tepoxalin. **a**, ABCB1 western blot with and without CRISPR knockout of ABCB1 in the LS1034 colon cancer cell line. **b**, Percent indel formation at genomic cut site in LS1034 ABCB1 CRISPR knockout lines assessed by NGS CRISPR assay. **c**, Cellular viability of LS1034 CRISPR knockout lines after treatment with tepoxalin for 5 days. Standard deviation across three replicates is shown with error bars. **d**, ABCB1 western blot with and without overexpression of ABCB1 in the Kuramochi ovarian cancer cell line. **e**, Cellular viability of Kuramochi wild type and ABCB1-overexpressing cells after treatment with tepoxalin for 8 days. Standard deviation across four replicates is shown with error bars. **f**, (Left) Single-agent dose-response curves for LS1034 cells treated with tepoxalin and zosuquidar for 5 days. Two replicates were averaged. (Middle) Dose response curves for tepoxalin in combination with varying doses of zosuquidar (indicated by different colors) for 5 days. Data are shown for tepoxalin doses above 470nM (the range indicated by the vertical dashed lines above). (Right) Bliss synergy scores estimated for each dose combination, showing strong antagonism by zosuquidar at tepoxalin doses above ∼5 μM. Two combinations were not tested (NT). **g**, Maximum synergy score is shown across several drug combinations measured in both LS1034 and REC1 cell lines. Both tariquidar and elacridar were strongly synergistic in combination with paclitaxel, while all MDR1 inhibitors tested were antagonistic in combination with tepoxalin.

**Extended Data Fig. 13.**
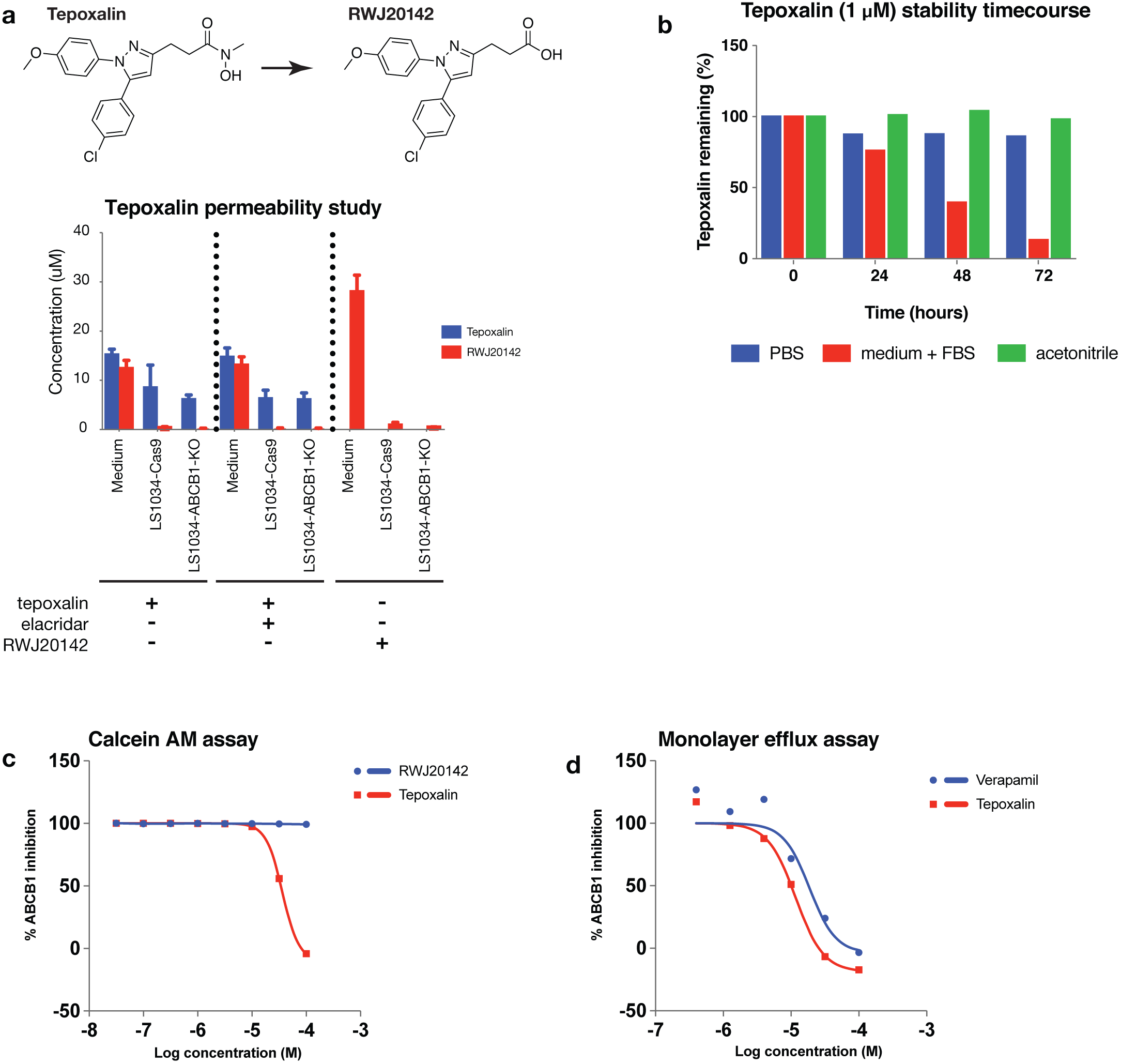
Tepoxalin is cell permeable and interacts with ABCB1 at high concentrations. **a**, Cellular permeability study of tepoxalin and RWJ20142 in LS1034 colon cancer cell lines with and without ABCB1 knockout. Each indicated cell line or media-only control was treated with 20 μM tepoxalin or RWJ20142 for 3 hours. Tepoxalin and RWJ20142 concentrations in cell lysate or medium were determined by liquid chromatography-mass spectrometry. Standard deviation is shown by error bars. **b**, Stability study of tepoxalin in three different vehicles over 72 hours. Percent of tepoxalin remaining is indicated at each timepoint. **c**, ABCB1 antagonism assay using calcein AM fluorescence. MDCKII cells were treated with indicated concentration of tepoxalin or RWJ20142. Mean percentage of ABCB1 inhibition across three replicates is shown. **d**, ABCB1 transport-based activity assay. Basolateral transport of the ABCB1 substrate loperimide was assessed using a monolayer of MDCK-ABCB1 cells in the presence of tepoxalin or RWJ20142. Mean percentage of ABCB1 inhibition across two replicates is shown.

